# Land use change through the lens of macroecology: insights from Azorean arthropods and the Maximum Entropy Theory of Ecology

**DOI:** 10.1101/2021.09.14.460355

**Authors:** Micah Brush, Thomas J. Matthews, Paulo A.V. Borges, John Harte

## Abstract

Human activity and land management practices, in particular land use change, have resulted in the global loss of biodiversity. These types of disturbance affect the shape of macroecological patterns, and therefore analyzing these patterns can provide insights into how ecosystems are affected by land use change. We here use arthropod census data from 96 sites at Terceira Island in the Azores archipelago across four different land uses of increasing management intensity: native forest, exotic forest, semi-natural pasture, and intensive pasture, to examine the effects of land use type on three macroecological patterns: the species abundance distribution, the metabolic rate distribution of individuals, and the species–area relationship. The Maximum Entropy Theory of Ecology (METE) has successfully predicted these patterns across habitats and taxa in undisturbed ecosystems, and thus provides a null expectation for their shapes. Across these patterns, we find that the forest habitats are the best fit by METE predictions, while the semi-natural pasture is consistently the worst fit, and the intensive pasture is intermediately well fit. We show that the direction of failure of the METE predictions at the pasture sites is likely due to the hyper-dominance of introduced spider species present there. We hypothesize that the particularly poor fit for the semi-natural pasture is due to the mix of arthropod communities out of equilibrium, leading to greater heterogeneity in composition and complex dynamics that violate METE’s assumption of static state variables. The comparative better fit for the intensive pasture plausibly results from more homogeneous arthropod communities that are well adapted to intensive management, and thus whose state variables are less in flux. Analyzing deviations from theoretical predictions across land use type provides useful information about how land use and disturbance affect ecosystems, and such comparisons could be useful across other habitats and taxa.

## 1 Introduction

Land management and altered land use is a primary driver of ecological disturbance worldwide (Foley et al. 2005; Pereira et al. 2012; Klein Goldewijk et al. 2017). Land use changes affect landscape heterogeneity, and result in the broad scale loss and fragmentation of natural habitats, creating a mosaic of habitat types in many landscapes (Fahrig 2003; Fischer and Lindenmayer 2007; Cardoso et al. 2009; Fahrig 2019). This type of human driven disturbance has resulted in global biodiversity loss (Martins et al. 2014; Pimm et al. 2014; Newbold et al. 2015; Maxwell et al. 2016; Newbold et al. 2018). On oceanic islands, the conversion of native vegetation to forestry (managed forest plantations – monocultures of fast growing trees), agricultural, and pasture land has had particularly severe impacts on the native biota due to the small-scale nature of islands, the sensitivity and small ranges of many island endemics, and the fact that land-use change on islands has often been accompanied by the spread of exotic species (Gillespie and Roderick 2002; Borges et al. 2006; Whittaker and Fernández-Palacios 2007; Gillespie et al. 2008; Whittaker et al. 2017).

Anthropogenic disturbance impacts the form of macroecological patterns (Gray et al. 1979; Hill and Hamer 1998; Dornelas et al. 2009; Newman 2019), including through land use changes (Simons et al. 2015; Xu et al. 2019). An effective method for analyzing the impacts of disturbance on biodiversity is comparing empirical patterns to theoretically expected shapes (Kempton and Taylor 1974; Carey et al. 2006; Supp et al. 2012; Matthews and Whittaker 2015; Newman et al. 2020; Franzman et al. 2021) and relating deviations to the type of disturbance. To interpret macroecological patterns in this way, we require a theoretical prediction for what these different patterns should look like in ecosystems that have not been disturbed or managed.

In this study, we use the Maximum Entropy Theory of Ecology (METE) for our theoretical predictions (Harte et al. 2008; Harte 2011; Harte and Newman 2014; Brummer and Newman 2019). METE has the advantage of simultaneously predicting many macroecological patterns and has been found to well describe empirical patterns across diverse taxa and habitats (Harte 2011; White et al. 2012; Xiao et al. 2015). Additionally, there is increasing evidence that METE predictions perform less well in disturbed ecosystems (Carey et al. 2006; Rominger et al. 2016; Newman et al. 2020; Franzman et al. 2021; Harte et al. 2021), which supports its use as a null theory. METE uses the principle of maximizing information entropy, and is characterized by three so-called state variables that constrain the predicted distributions for a given ecosystem or habitat: the species richness *S*_0_, the number of individuals *N*_0_ and the total metabolic rate *E*_0_. To make spatial predictions, METE also requires the total area of the site *A*_0_.

The effects of land use change on the deviation of macroecological patterns from METE predictions has not yet been explored, as most disturbance has been characterized by ecosystems with rapidly changing state variables. Given that METE predictions appear to perform better in pristine ecosystems where state variables are changing relatively slowly in time, we expect that anthropogenic land uses that introduce significant disturbance should result in patterns that deviate from the predictions in meaningful ways (Harte et al. 2021). How well the data fit METE across land use types can thus provide insights about how different land uses affect these large scale patterns, and by extension how land use change affects biodiversity.

Here, we investigate how land use change affects several patterns predicted by METE with arthropod data from Terceira Island in the Azores archipelago (Portugal). The Azores are an isolated island chain in the Atlantic Ocean that have been populated for about 600 years (Norder et al. 2020) and have undergone extensive land use change since human colonization of the islands Previous work based on the analysis of Azorean arthropod data has shown that a variety of macroecological patterns, such as the species abundance distribution and functional trait composition, vary as a function of land use (Fattorini et al. 2016; Borda-de-Água et al. 2017; Rigal et al. 2018). Thus, it represents an ideal system to test how land use changes affect macroecological patterns and their deviation from METE predictions.

We test three predictions of METE simultaneously: the species abundance distribution (SAD), the metabolic energy rate distribution of individuals (MRDI), and the species–area relationship (SAR). See Table 1 for more information about these patterns and their METE predicted forms. Analyzing multiple patterns simultaneously avoids treating any individual pattern in isolation, especially since single patterns can often be predicted from many different underlying theories (McGill et al. 2007). We compare all three of these patterns using arthropod data across four land use types on Terceira Island, and analyze the deviations from the predicted patterns for information about how disturbance is affecting species community assembly. We predict that METE predictions will fit the data better for less intensively managed land uses, with deviations linearly increasing with management intensity.

**Table 1:**
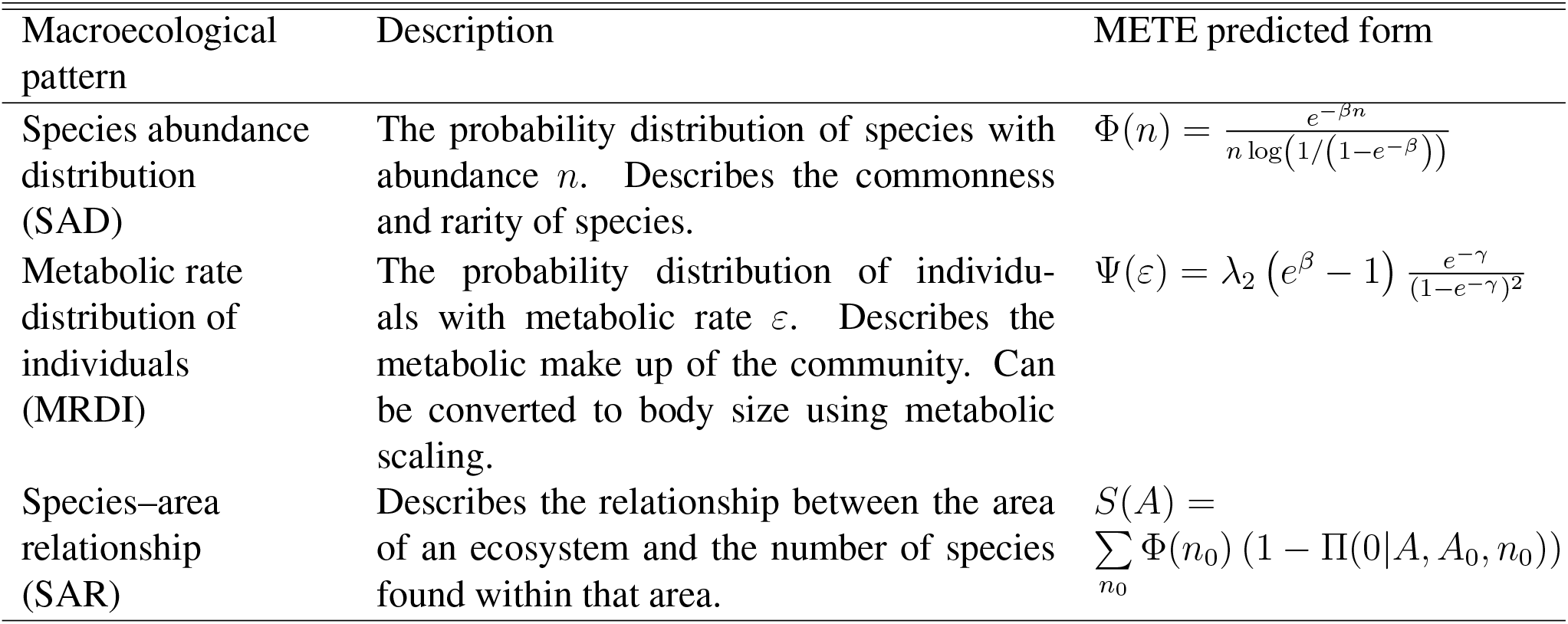
Descriptions of the macroecological patterns used in this study. The forms predicted by the Maximum Entropy Theory of Ecology are shown in the right column, where *β* = *λ*_1_ + *λ*_2_, and *λ* = *λ*_1_ + *λ*_2_*ε*, and *λ*_1_ and *λ*_2_ are calculated from the species richness *S*_0_, number of individuals *N*_0_, and total metabolic rate *E*_0_ (see Appendix S3). The species–area relationship must be calculated at each new scale *A* from an existing scale *A*_0_ using the species abundance distribution Φ(*n*) and the species-level spatial abundance distribution Π(*n*).

Additionally, many arthropod species have been introduced to the archipelago by humans (Borges et al. 2010). These exotic species have changed the ecological landscape (Florencio et al. 2013) and have been found to have a different functional trait composition from the indigenous species (Rigal et al. 2018). However, some studies have found that these species appear to be integrated in these ecosystems, perhaps by replacing lost indigenous species and/or filling empty niche space (Gaston et al. 2006; Rigal et al. 2013). We might therefore expect that METE predictions would perform better when indigenous and introduced species are analyzed together. To test this, we analyze indigenous and exotic species separately, in addition to our analyses with all species together.

## 2 Methods

### 2.1 Study area and arthropod data

The Azores Islands are an isolated island chain in the Atlantic Ocean of volcanic origin. All of the data analyzed here come from Terceira Island, which before human colonization was almost entirely forested but now comprises a mix of land uses. The four major land uses, ranked in increasing order of management intensity, are 1. native forest, 2. exotic forest, 3. semi-natural pasture, and 4. intensive pasture (Cardoso et al. 2009; Rigal et al. 2018). These land uses comprise about 87% of the total island area, which is broken down by land use in Table 2 (Cardoso et al. 2009). Figure S1.1 in Rigal et al. (2018) shows a land use distribution map of Terceira Island with more specific spatial information.

**Table 2:** The total number of species and individuals observed for each land use, and the median number across sites within one land use. The number in parentheses is the number of indigenous species or individuals, followed by the number of exotic species or individuals. Additionally, the number of sites where data were collected for each land use, and the percent of the total island area occupied by that land use. Across all land uses, there are a total of 271 species and 46 250 individuals, with four species constituting 11 individuals that are not identified as indigenous or exotic. The dataset in full is used for the SAD analysis, but for the MRDI analysis juvenile individuals were excluded, leaving a total of 226 species and 36 269 individuals, and for the SAR analysis there are less data available with the spatial resolution needed, leaving 228 species and 34 282 individuals.

| Land use | % Area | Sites | Total $S_0$ | Total $N_0$ | Median $S_0$ | Median $N_0$ |
| --- | --- | --- | --- | --- | --- | --- |
| Native forest | 9 | 44 | 148 (86, 60) | 10 291 (7288, 3001) | 24 (16, 8) | 195 (129, 50) |
| Exotic forest | 15 | 12 | 87 (44, 42) | 3385 (1476, 1908) | 20 (10, 9) | 196 ( 11, 51) |
| Semi-natural pasture | 15 | 16 | 127 (50, 76) | 11 421 (2110, 9310) | 28 (10, 17) | 766 (101, 623) |
| Intensive pasture | 48 | 24 | 136 (40, 94) | 21 153 (4076, 17 070) | 36 (10, 27) | 878 (161, 684) |

The native forest is made up of perennial trees and shrubs adapted to a hyper-humid Atlantic climate, and is now restricted to elevations above 500 m above sea level (a.s.l.) and dominated by *Juniperus-Ilex* forests and *Juniperus* woodlands (Elias et al. 2016). Exotic plantations of the fast growing tree *Cryptomeria japonica* were planted after the Second World War to reforest large areas of previous native forest that were destroyed in the previous decades for fuel. These plantations are dense and almost no understory is present. Semi-natural pastures are located around 400-600 m a.s.l., have a mixture of native and exotic herbs and grasses, and are mostly grazed in the spring and summer with low cattle density. Intensive pastures are located between 100-500 m a.s.l. and are grazed every three weeks (and sometimes up to every 12 days in the summer) with high cattle density.

The arthropod samples were collected using pitfall traps across 96 sites. Each of the sites has a single 150 m transect with 30 pitfall traps spaced out at 5 m intervals: 15 traps filled with approximately 60 mL of a non-attractive solution (anti-freeze liquid) with a small proportion of ethylene glycol, and 15 traps with the same volume of a general attractive solution (Turquin), which was made of dark beer and some preservatives. All data were collected over summers on Terceira Island over the period from 1997-2009 (for more details, see Borges et al. 2005; Cardoso et al. 2009; Rigal et al. 2018).

Table 2 shows the number of sites for each land use, along with the total and median number of species *S*_0_ and individuals *N*_0_ across all transects in that land use (including indigenous and exotic). Indigenous species are those that are endemic (occur only in the Azores) or native (appear in the Azores Islands and other nearby archipelagos and/or the mainland). Exotic, or introduced, species are those believed to have been introduced by humans following human colonization of the archipelago in the 15th century (Borges et al. 2010). Unidentified species that share a genus, subfamily, or family with other species present in the archipelago are put into the same colonization category as those species (Borges et al. 2010; Florencio et al. 2013). Four remaining species (11 individuals) that are not identified as indigenous or exotic are removed from the analysis when indigenous and introduced species are analyzed separately. Across all land uses, there are a total of 271 species and 46 250 individuals, with 126 indigenous species and 14 950 indigenous individuals and 141 exotic species and 31 289 exotic individuals. However, juvenile individuals are excluded from the MRDI analysis, leaving 226 species and 36 269 individuals, and there are less data available with the spatial resolution needed for the SAR analysis, with 228 species and 34 282 individuals.

Body length measurements of individuals from the different species are obtained as described in Rigal et al. (2018), and average body length values are used here. For 26 Araneae species, we use updated body length measurements taken from a new database of Macaronesian spider traits (Macías-Hernández et al. 2020). Body lengths are then converted to body mass values using empirical scaling equations, as detailed in Appendix S1 (Hódar 1996; Baumgärtner and Rothhaupt 2003; Wardhaugh 2013). Additionally, body mass variation within a single species is reintroduced by assuming a normal distribution for intraspecific body size (Gouws et al. 2011), and then obtaining parameters for this distribution by relating the mean and variance of body mass for several spider (Macías-Hernández et al. 2020) and beetle (Terzopoulou et al. 2015) species. We then simulate body masses for all individuals using the parameters obtained for beetles for all orders except for spiders, where we use the parameters obtained for them. Since Coleoptera and Araneae are the two most common orders in the dataset, differences among other orders should not overly impact the analysis. For more information on how we introduce intraspecific body mass variation, see Appendix S2. We then use metabolic scaling to convert body mass data to metabolic rate, assuming that *ε* α *m*^3/4^, where *m* is the body mass, and rescaling such that the smallest *ε* = 1.

### 2.2 Comparing METE predictions with data

We divide the data by land use and compare the observations to the predictions of METE, which are reviewed in Appendix S3 and summarized in Table 1. We primarily analyze our results at the individual transect level, treating each transect in one land use category as a replicate. This is because METE makes predictions within a single community, and the number of species across many small patches is not the same compared to a single large patch of the same area. However, we find mostly similar results when combining data from all transects for each land use (Appendix S4).

Our sampled arthropod data are separated into juvenile and adult individuals. For the SAD and SAR analysis, we treat these together as a single dataset that accurately captures all ground dwelling arthropods. For the MRDI, the empirical metabolic rates are calculated from scaling relationships using body length data which were only available for adult arthropods (see Section 2.1 and Appendix S1), and thus we were unable to include the juvenile individuals in our analysis.

For the SAD and MRDI, we use the mean least squares of the log of the abundance or metabolic rate as our primary goodness of fit metric. Mathematically, for the SAD, this means we take 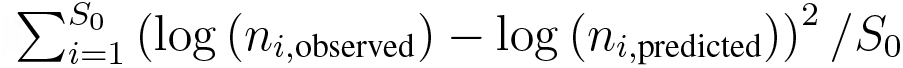, where *n*_*i*_ is the abundance of the species with rank *i*, and we take the mean over all *S*_0_ ranks. For the MRDI, we replace *n* with *ε*, and *S*_0_ with *N*_0_ as the distribution is over individuals rather than species. To ensure our results are robust to our choice of goodness of fit metric, we additionally perform Kolmogorov-Smirnov goodness of fit tests for the empirical CDF compared to the METE predicted CDF for both the SAD and MRDI and obtain similar results to the mean least squared analysis (Appendix S5).

For the SAR, we compare the predicted and empirical number of species across scales of 1, 2, 3, 5, 6, 10, 15, and 30 traps which are evenly spaced at 5m intervals (Section 2.1). These relative scales were chosen as they use all data available at each scale. Given that METE predicts that all nested SARs will collapse onto a single universal curve when plotted as the slope of the SAR *z* versus *D* = log(*N*_0_/*S*_0_), a scale parameter (Harte 2011; Wilber et al. 2015), we primarily compare the predicted and observed slopes at each scale (though we find similar results comparing the number of species at each scale directly, see Appendix S6). Given the number of species and individuals at a given scale, we predict the number of species at the next smallest scale and then calculate the corresponding slope. Mathematically,

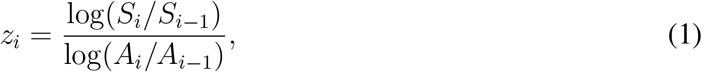

where *i* indexes the scale. In the list of scales above, *i* = 1 corresponds to the average number of species at the scale of 1 cell, and *i* = 8 corresponds to the number of species in all 30 cells. The empirical slope is obtained by comparing the average number of species at the scale under consideration to the average number of species at the next smallest scale, as in Eq. 1, but now the smaller scale is also empirical. We then compare slopes at all but the smallest scale, leaving seven scales of comparison, and again take the mean least squares 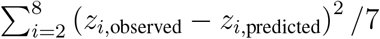, where *i* indexes the scale and we do not compare at the smallest scale *i* = 1. We additionally only use scales where the empirical average for the number of species is greater than four (*S*_0_ > 4), and so for many transects we will have fewer than seven data points. This is because several METE simplifications break down for small *S*_0_.

For all patterns, we ran the analyses using all species, and exotic and indigenous species separately. Additional details on how we compared METE predictions to data can be found in Appendix S7.

Finally, we analyzed the differences in the mean least squared error results across land uses for each pattern when all species were combined. We used the ANOVA *F*-test in combination with the Tukey post-hoc test, as well as the Kruskal-Wallis test in combination with the Dunn post-hoc test (see Appendix S8).

## 3 Results

We first present the results for each pattern when all species are combined, followed by a separate section with results relating to the analysis of the indigenous and introduced species separately. Fig. 1 shows the mean deviation from METE across land use type and pattern. The markers in Fig. 1a show the mean and standard error of the distribution of mean least squared error over transects at each land use for each of the three macroecological patterns with all species combined, and Fig. 1b shows the same results when indigenous and introduced species are considered separately. Statistical analyses (the ANOVA *F*-test, the Tukey post-hoc test, the Kruskal-Wallis test, and the Dunn post-hoc test) robustly support the differences in deviation between land uses that can be seen in Fig. 1a (Appendix S8), and our results are generally consistent when analyzed at the community level (Appendix S4) and when using the KS test statistic rather than mean least squared error (Appendix S5).

**Figure 1:**
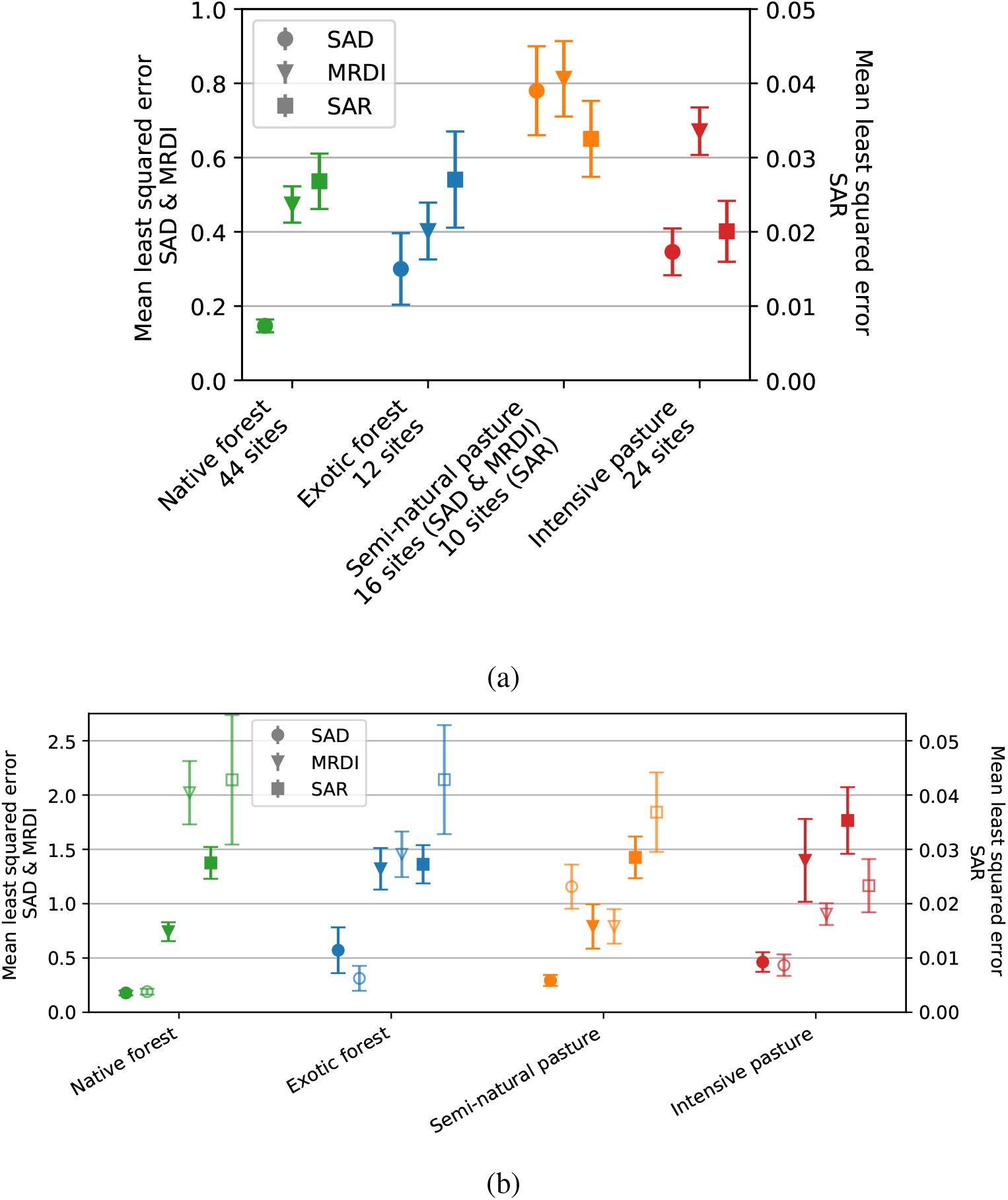
Mean and standard error of the distribution of mean least squared errors calculated from each transect, across all three patterns (SAD, MRDI, SAR) and land use types. Fig. 1a shows the results when all species are analyzed together, and Fig. 1b shows the results for indigenous (filled shapes) and introduced species (open shapes, lighter color) separately. For the SAD and the MRDI, the mean least squared error is the mean of the squared difference between the observed values of the rank ordered natural log of the abundance or metabolic rate, respectively, minus the METE predicted values. For the SAR, the mean least squared error is the mean of the squared observed minus the predicted value for the slope, *z* across scales. Note the difference in y-scale for the SAR, where the mean least squared error was much smaller. The shape of the marker indicates the pattern and the color indicates the land use. For Fig. 1b, the number of sites for the SAD are 44, 12, 16, and 24, for the MRDI (44,32), (11,12), (15,16), and (24,24), and for the SAR (43,9), (10,9), (9,10), and (20,24), in order of increasing land use intensity, and where the first number in parentheses is the number of sites with indigenous species and the second with introduced.

### 3.1 Species abundance distribution (SAD)

We find that the semi-natural pasture is particularly poorly described by METE, and the native forest is the best fit, with the results for the exotic forest and intensive pasture being fairly similar and intermediate (Fig. 1a, Table S2). The standard error of the mean is the lowest for the native forest sites, and the highest for the semi-natural pasture. Similar results for the KS test statistic *D*_*KS*_ across transects are shown in Appendix S5.

To combine all of the SADs for each land use onto a single plot, we plot the residuals of log_10_(abundance) in Fig. 2. The residuals are calculated as log_10_(abundance_observed_) - log_10_(abundance_predicted_), and the x-axis has been scaled by the number of species to facilitate comparison between sites with different total number of species. Therefore, each line in this plot represents the deviation of the SAD from the METE prediction at a single site. For clarity, the null expectations for the residuals are shown in Appendix S9. Additionally, plots of the SAD for each transect are shown in Appendix S10.

**Figure 2:**
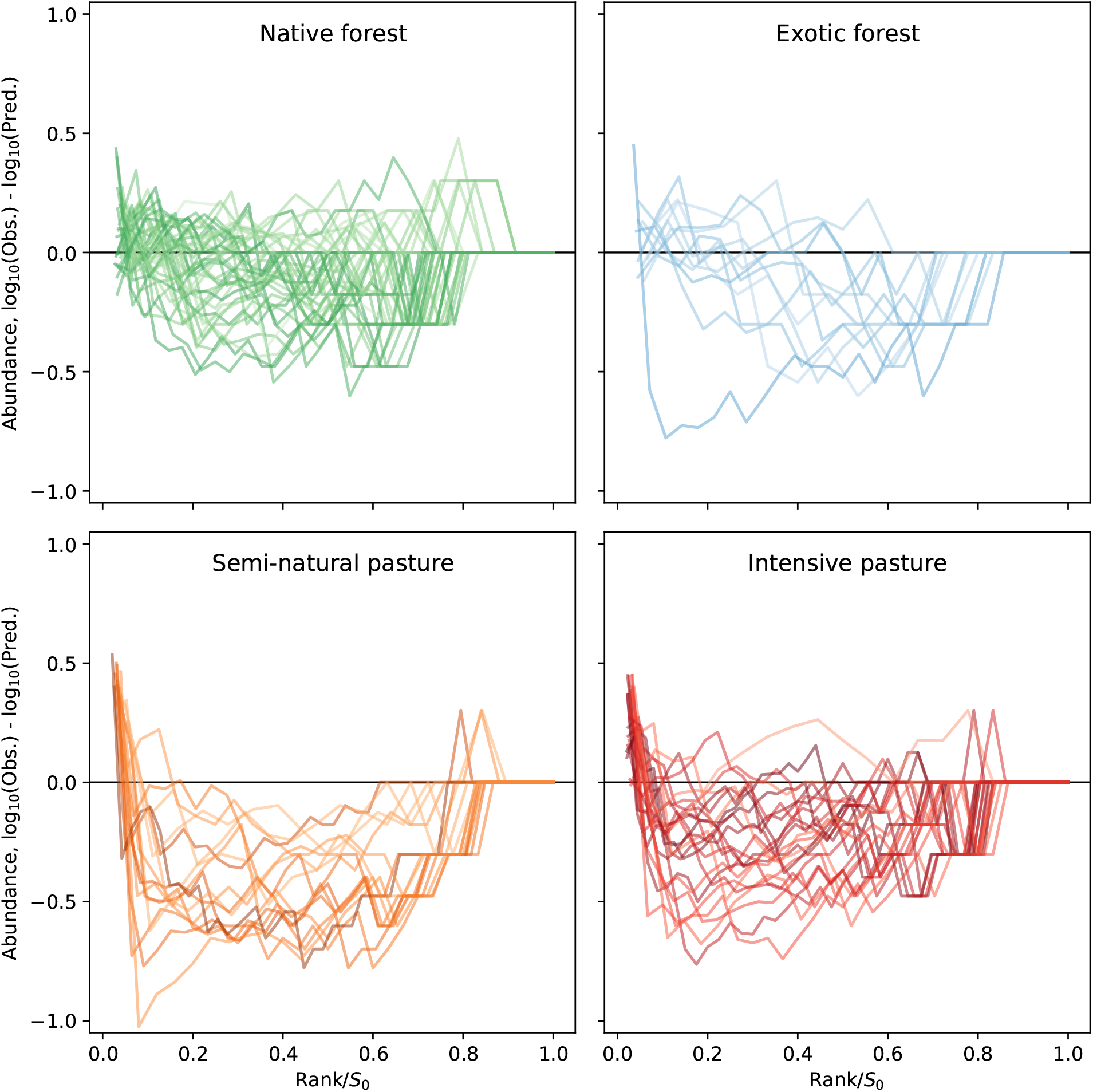
The residuals of log_10_ of the observed abundance minus log_10_ of the predicted abundance from METE for each transect across land uses. As the number of ranks is equal to the number of species *S*_0_, the ranks on the x-axis have been rescaled by 1/*S*_0_ to facilitate comparison between sites. The darker lines are sites with a higher number of species, and lighter lines represent sites with fewer species. The colors correspond to the different land uses.

In the semi-natural pasture sites, we see that METE consistently under predicts the abundance of the most abundant species, as the residuals are consistently well above the zero line at low rank across all transects. This is in contrast to the null expectation that the most abundant species across transects should be occasionally under and over predicted, if the abundances were drawn randomly from the METE predicted log series distribution. METE additionally over predicts the abundance of the species of intermediate rank, as the residuals dip below the zero line at intermediate rank for all transects in this land use. Again here, the null expectation would be that the residuals should be scattered both above and below the zero line across rank. We can see a similar pattern across land uses where the residuals of the most abundant species tend to be positive, though it is most prevalent at the pasture sites and least common at the native forest sites. Across all sites, METE generally under predicts the number of singletons, though this is again less common in the native forest sites.

Results where we have combined all transects together rather than treating them as replicates at each land use can be found in Appendix S4. In that case, the forest sites are again better fit than the pasture sites, but the ranking is slightly different.

### 3.2 Metabolic rate distribution of individuals (MRDI)

The forest sites are better fit by METE than the pasture sites, and the difference between the forest sites is small (Fig. 1a, Table S3).

As with the SAD we plot the residuals for each transect, calculated as observed minus predicted of log_10_ of the metabolic rate, in Fig. 3, where this time the x-axis has been scaled by the number of individuals to facilitate comparison across transects. Again, each line here represents a single site at that land use and indicates how that transect deviates from the METE prediction. If the transects were randomly sampled from the METE distribution we would expect random scatter above and below the zero line in this plot, which is not observed as there are clear patterns of deviation. The null expectations for these residuals are shown in Appendix S9. The rank ordered plots for the MRDIs at each transect are plotted in Appendix S11.

**Figure 3:**
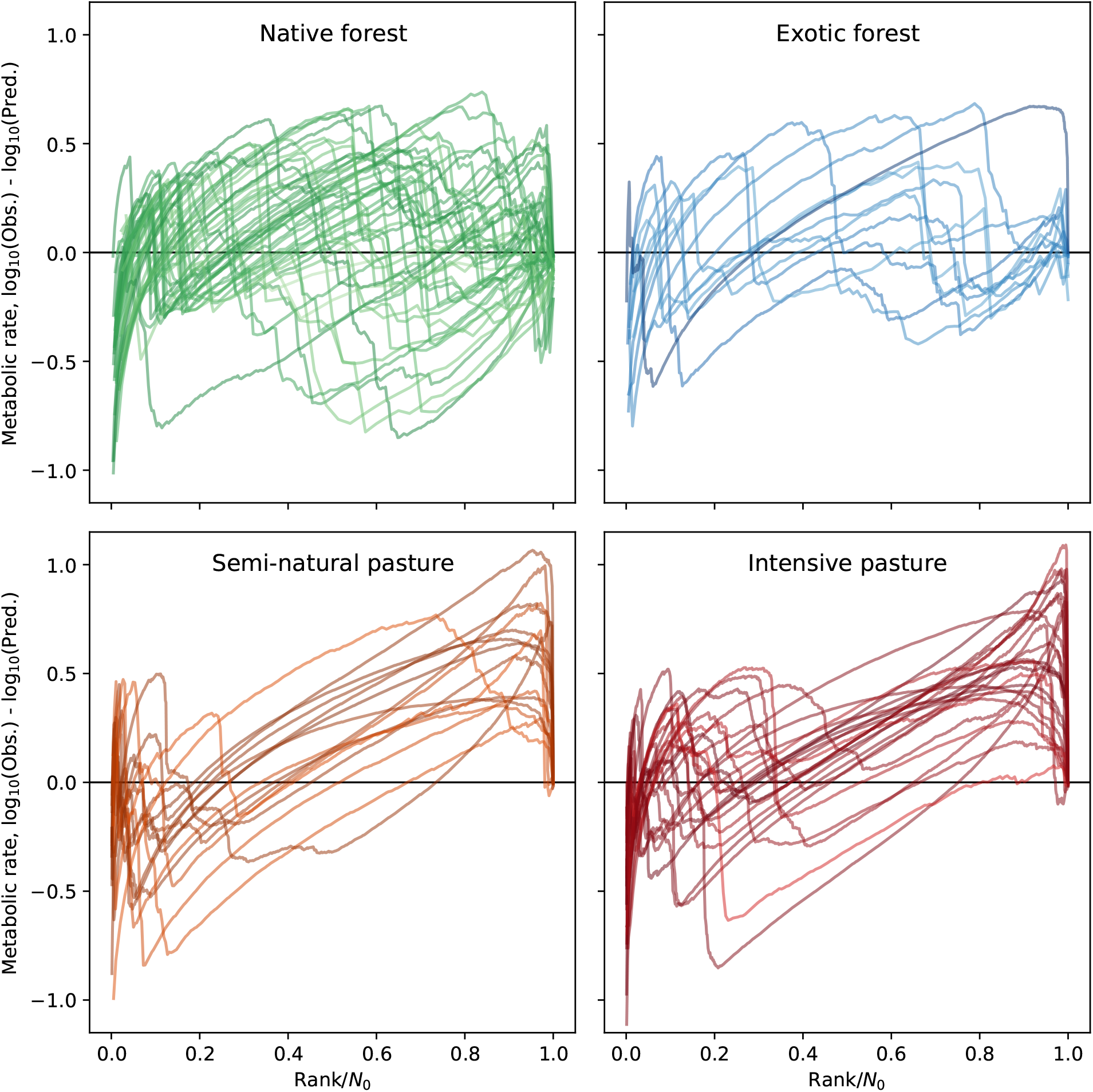
The residuals of log_10_ of the observed metabolic rate minus log_10_ of the predicted metabolic rate for the rank ordered plots. As the number of ranks is equal to the number of individuals *N*_0_, the ranks on the x-axis have been rescaled by 1/*N*_0_ to facilitate comparison between sites. The darker lines are sites with a higher number of individuals, and lighter lines represent sites with fewer individuals. The colors correspond to the different land uses.

Across land uses in the residuals, we see long lines of constant slope, particularly at the pasture sites. This pattern appears because METE predicts that the metabolic rate should be proportional to 1/rank for intermediate to large rank (Table 1), but we observe many individuals of similar metabolic rate at these ranks because the highly abundant species are small bodied and therefore have low metabolic rate (for specific examples, see many of the individual transects in Appendix S11). This means that METE initially overpredicts the metabolic rate of these species, but because there are so many individuals at a similar metabolic rate, and the METE prediction decays rapidly, METE underpredicts the metabolic rate at higher ranks. This leads to the long lines of near constant slope observed in Fig. 3, which does not match the null expectation of scatter around the zero line.

The results are again similar if analyzed using the KS test statistic (Appendix S5) or at the community level (Appendix S4, though here the fit for the intensive pasture is much worse).

### 3.3 Species–area relationship (SAR)

The mean deviations from METE for the SAR are comparable across land use types, though the semi-natural pasture does appear to give the worse fit. Note the different y-axis scale in Fig. 1a compared to the SAD and MRDI, and that the mean least squared error for each transect is averaged over the number of scales where the empirical *S*_0_ > 4.

Figure 4 compares the SAR data for each site, organized by land use. Each point here represents a single transect at a single scale *D* = log (*N*_0_/*S*_0_), and the lines are the corresponding METE predictions. There is a large amount of scatter in these plots across land use types. We see that METE tends to under predict the slope at larger scales, and thus there is bias in the direction of the deviation from METE. Section Appendix S12 (Fig. S19) uses the scale-collapse of the *z* − *D* relationship to display all of the data across sites on one plot.

**Figure 4:**
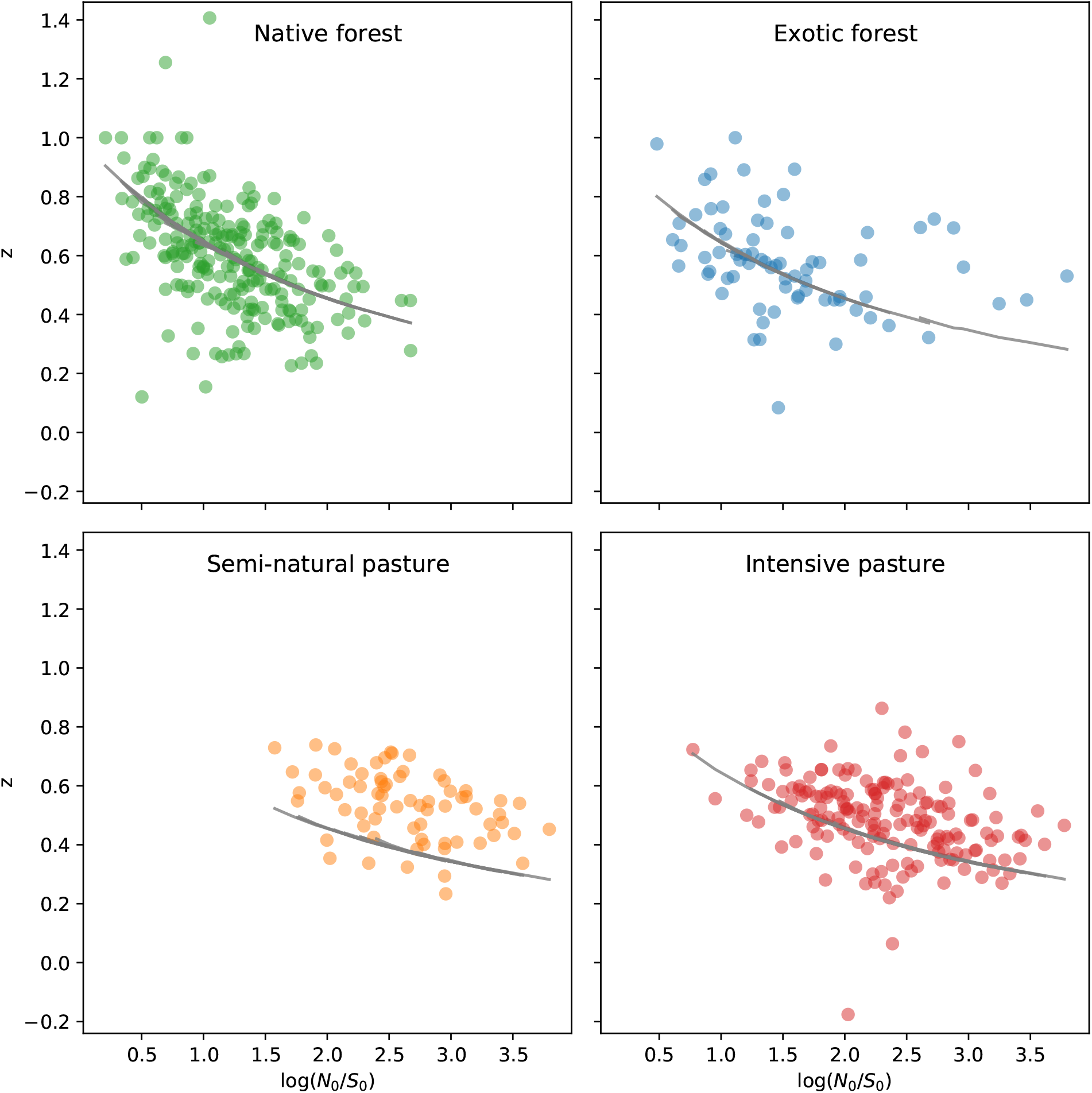
The species–area relationship for each transect across land use. Each point represents a single transect at a specific scale, where the scale is determined by *D* = log(*N*_0_/*S*_0_), and are colored according to land use type. The gray lines are the METE predictions, which largely overlap due to the scale collapse prediction of METE. Here we have plotted the slope of the relationship on the y-axis so that all points collapse onto one universal curve.

The results are similar if we analyze the predicted number of species at each scale rather than the slope (Appendix S6).

### 3.4 Indigenous and exotic species

Figure 1b shows the mean and associated standard error of mean least squares across transects, separated by species that are indigenous and introduced, at all four different land uses. The number of sites where we are able to separate the individuals in this way is limited for the MRDI as adults of all species are not present at each site, and for the SAR as we again only use scales where *S*_0_ > 4. Note the difference in y-axis scales between Fig. 1a and Fig. 1b, which shows that the error when the indigenous and introduced species are separated is generally larger than when the species are considered together.

For the SAD, the largest difference between the indigenous and introduced species is at the semi-natural pasture sites, where the introduced species fit quite poorly compared to the indigenous species. Across other land uses indigenous and introduced species are comparably well fit, though the introduced species fit slightly better at the exotic forest sites. These fits are also generally comparable to the combined fits, except that the fit of the indigenous species at the exotic forest and the introduced species at the semi-natural pasture are worse (Fig. 1a).

For the MRDI, we find a large difference at the native forest sites, where the introduced species again fit poorly compared to the indigenous species. We see a slight difference at the intensive pasture site, where the introduced species fit slightly better. We also find a very different trend overall, in that the semi-natural pasture is no longer the worst fit to the METE predictions. For the indigenous species the goodness of fit is similar between the native forest and the semi-natural pasture, and is worse but again similar at the exotic forest and intensive pasture sites. For the introduced species, the semi-natural pasture has the best fit, though the intensive pasture is close, and at both forest sites the fit is much worse than when the species are analyzed together. Additionally, the fits for the introduced species in the native forest and the introduced and indigenous species at the exotic forest are all worse than the fit when the species are combined (Fig. 1a).

For the SAR, the larger error and smaller differences make the analysis more difficult, though we do seem to see a trend of decreasing mean least squares with land use intensity for the introduced species, and increasing mean least squares for the indigenous species. This means that for the first three land uses, the indigenous species are better fit, and only at the intensive pasture are the introduced species better fit. The fit for the indigenous species across all land uses except the intensive pasture is comparable to that of the combined fit (Fig. 1a).

## 4 Discussion

In this study we have compared, for three commonly studied macroecological patterns (the SAD, MRDI, and SAR), the predictions of METE to the empirical patterns of Azorean arthropods across four land use types of varying management intensity. Overall we find that when all species (indigenous and exotic) are considered together, METE provides a reasonable approximation to the patterns in the native forest and exotic forest sites, as predicted. In contrast, METE provides the worst fit to the semi-natural pasture sites. Interestingly, METE provides a relatively better fit, similar to that for the exotic forest sites, to the most intensively managed land use sites, the intensive pasture. These conclusions can be seen in Fig. 1a as differences between the mean deviation from METE across land use and pattern, and are supported by the statistical analyses in Appendix S8. We now discuss the results for each of the three patterns in turn. Unless otherwise noted, we discuss the results relating to the analyses where the indigenous and introduced species are combined.

### 4.1 Species abundance distribution (SAD)

The most distinctive pattern in the SAD residuals in Fig. 2 is the consistent under prediction of the most abundant species, particularly at the pasture sites. This result is comparable to the findings of Simons et al. (2015) and Xu et al. (2019), who found that increased grazing (Xu et al. 2019) or land use (Simons et al. 2015) intensity led to a reduction in evenness and an increase in the dominance of the most abundant species (ie. hyper-dominance sensu Hubbell 2013). We attribute the dominance of a few very abundant species to small-bodied, highly dispersive, mostly introduced spider species, which were found to be very prevalent at sites with high land use intensity (Borges and Wunderlich 2008; Rigal et al. 2018). Here, we find that the most abundant species in both the semi-natural pasture and the intensive pasture is *Oedothorax fuscus*, which is indeed a small-bodied introduced spider that lives mostly in grasslands. This spider accounts for 67% of all individuals in the semi-natural pasture, and 43% of all individuals in the intensive pasture. Another example of this type of spider is *Erigone dentipalpis*, which is the fourth most common species at the semi-natural pasture sites and the second most common species at the intensive pasture sites, accounting for 2% and 9% of all individuals, respectively. These *Erigoninae* spiders, being highly dispersive, are adapted to the disturbance caused by the grazing performed every three weeks in the intensive pastures and the more intensive grazing performed in semi-natural pastures during summer.

A previous study using Azorean arthropod data found that species dispersal ability affects the form of the SAD (Borda-de-Água et al. 2017). It was found that when species are grouped together by dispersal ability, the corresponding Preston plots (number of species versus log_2_(abundance)) are steep for high dispersal species and develop an intermediate mode for lower dispersal species. When rank ordered, this results in SADs that are steeper at low rank for high dispersal species, and less steep at low rank for lower dispersal species. We see very steep rank ordered SADs in the pasture sites in Fig. 2 and in Appendix S10, which we postulate is a result of the highly dispersive spider species. These species are mostly introduced, and thus have not yet evolved to have reduced dispersal ability as they have only been present on the island for a short period of time relative to evolutionary time scales (Borges and Wunderlich 2008).

These mostly introduced spider species additionally have multiple generations per year, which allows them to recolonize the pasture sites from the nearby surrounding forest after disturbance due to land management or cattle grazing (see Rigal et al. 2018, Figure S1.1 for a map). This recolonization is particularly relevant for the semi-natural pasture sites, where cattle grazing and fertilization is seasonal (Borges and Brown 2004). The particularly high abundance of these spiders species in these data may also be related to the fact that these data were collected in the summers, when the semi-natural pasture sites are likely to be subject to cattle grazing.

METE also tends to under predict the number of singletons at most sites, except in some cases for the native forest sites where we see that METE over predicts the number of species with small abundance. This could be related to sampling, as the traps are less likely to capture multiple individuals of rare species, but it could also be related to the METE prediction for the number of singletons. METE predicts a number of singletons equal to *βN*_0_, and therefore increasing *N*_0_ while holding *S*_0_ constant decreases the expected number of singletons (Harte 2011, Chapter 7.3). From Table 2, we see that *N*_0_/*S*_0_ is large for the pasture sites compared to the forest sites, and therefore METE predicts proportionally fewer singletons at these sites. If ecologically we still expect a similar number of singletons, given that the high *N*_0_/*S*_0_ is being driven by a small number of very abundant species, then this could mean that METE under predicts the number of singletons.

### 4.2 Metabolic rate distribution of individuals (MRDI)

Across land use types, we find that the MRDIs are not particularly well described by the METE prediction. However, there are reasons we might expect this. For example, Xiao et al. (2015) discuss that we should not necessarily expect animals (rather than plants) to follow the METE predicted MRDI because animal body sizes for each species are likely to be clustered around an intermediate value (eg. Gouws et al. 2011), leading to multimodal MRDIs (Thibault et al. 2011) unlike the monotonically decreasing form predicted by METE. Our metabolic rate distributions are indeed multimodal at most sites. Additionally, we had to make a number of approximations to obtain these distributions, such as the use of scaling relationships and reintroducing intraspecies variation (see Appendix S1 and Appendix S2) Finally, only adults are included in the metabolic rate distribution, which could result in missing the lower end of the unscaled MRDI, resulting in a skewed distribution. As METE predicts relative metabolic rate (scaled so that the smallest organism is *ε* = 1 (Harte 2011)), this will result in over predicting the metabolic rate of the individuals with the greatest metabolic rate. Despite these issues, we can still quantify which land use is the most well described by METE, and we see a similar relationship between land use and goodness of fit when compared with the SAD and SAR.

At the pasture sites, we consistently under predict the low rank, high metabolic rate individuals, and over predict the high rank, low metabolic rate individuals, resulting in a pattern where the residuals have positive slope (Fig. 3). These patterns are indicative of a large number of individuals with similar metabolic rate (see Appendix S11 for plots of each site). As discussed in relation to the SAD, these sites have a few highly abundant, small bodied spider species. These species have comparatively low metabolic rate, and the variation in metabolic rate within a species is smaller than the variation across species. We therefore end up with long lines of positive slope in the residuals as the METE prediction slopes downward over rank but the empirical MRDI remains roughly constant. We see this especially at intermediate and low ranks as these species have low metabolic rates. Thus, this pattern is also likely driven by a few highly abundant species.

### 4.3 Species–area relationship (SAR)

The mean least squares comparisons for the SAR in Fig. 1a and Fig. S7 are noticeably different from those for the SAD and MRDI. Again, here we find that the semi-natural pasture is the worst fit by METE, but it is not as dramatic as in the other cases and the fit is much closer to both forest sites. Additionally, we find that the intensive pasture is the best fit by METE, though it is comparable to other land use types given the error. However, the mean least squares is not the only goodness of fit metric. Particularly in the case of the pasture sites here, we see clear a clear pattern that METE under predicts *z* and correspondingly over predicts the number of species at small scales. This is in line with our analysis of the SAD at the pasture sites, in that these sites have more high abundance species compared to the METE prediction. Overall, even though the mean least squares metric is smaller at the intensive pasture sites, the direction of the difference is more biased.

In general, the pasture sites correspond to larger *N*_0_/*S*_0_ than the forest sites (see Fig. 4 and Table 2). When using *D* = log(*N*_0_/*S*_0_) as a scale variable, this means that the pastures are testing a different scale compared to the forest sites. We see this in Fig. 4 (and more easily in Fig. S19), where the METE prediction for *z* is noticeably lower than the data points starting around log(*N*_0_/*S*_0_) ≈ 2, which is also where most of the pasture data points are clustered. This could indicate that the failure of METE to accurately predict the SAR is coming more from the underlying prediction of a log series abundance distribution (Appendix S3), rather than from the species-level spatial abundance distribution prediction, as the log series prediction for the pasture sites under predicts the abundance of the most abundant species.

### 4.4 Indigenous and exotic species

The overall fit is largely comparable when indigenous and exotic species are considered together and separately, and in many cases there is little difference between the indigenous and introduced species (Fig. 1). This is in line with previous studies that found that exotic species were integrated with indigenous species in the Azorean arthropod communities by analyzing the interspecific abundance-occupancy relationship (Gaston et al. 2006; Rigal et al. 2013).

In terms of METE, we expect that the maximum entropy inference technique should apply to any collection of entities, however they are categorized. For example, using arthropod data from Panama, Harte and Kitzes (2015) found evidence that the analysis of the form of the SAD is relatively insensitive to the choice or taxonomic category used for the analysis. Our findings here provide further support for a flexible application of METE across colonization categories.

Given the similarity of trends between Fig. 1a and 1b, our discussion here is quite speculative. We note that for the SAD, the goodness of fit is comparable between indigenous and introduced species except at the semi-natural pasture sites, which supports our hypothesis that the poor fit there is driven by introduced species and could point to more complex dynamics between indigenous and introduced species at these sites (discussed in more depth below). For the MRDI and the SAR, but not for the SAD, the deviation from METE generally decreases with increasing land use intensity for the introduced species, which could be an expected trend if the introduced species are more adapted to those habitat types. For example, in the native forest, many of the human adapted introduced species are likely only present in a stochastic sense due to source-sink dynamics (Matthews et al. 2019; Borges et al. 2020), leading to poor fit with METE predictions.

### 4.5 Implications for future METE studies in anthropogenic landscapes

In other studies of METE, disturbance is often linked to rapid change in state variables (Newman et al. 2020; Franzman et al. 2021; Harte et al. 2021). The dynamics are then out of steady state, and the state variables alone are not sufficient to describe the macroecological patterns. Here, we instead analyze how land use change affects deviation from METE predictions assuming that the deviation in time at any given land use type is relatively static. Assuming that disturbance is connected to the rate of change of the state variables, we could interpret the poor fit of the semi-natural pasture as indicating that *N*_0_, *E*_0_ and/or *S*_0_ are not constant over ecological time scales. We could test this hypothesis with time resolved data of arthropod composition. For example, we might expect the state variables to change with management intensity over the year. It also may be the case that disturbance is more general and cannot always be characterized by changing state variables, and may depend on additional factors such as the rate of migration in and out of the ecosystem rather than just the net difference.

More broadly, we hypothesize that heterogeneity in species composition may lead to deviations from METE. It is possible that when species from different groups interact, their dynamics are more complicated and violate the underlying METE assumption that *N*_0_, *E*_0_, and *S*_0_ alone are adequate to characterize the larger scale patterns. These more complicated dynamics could, for example, be related to the mix of indigenous and introduced species, core and occasional species, or to source-sink dynamics (see Matthews 2021), depending on the habitat.

In this study, the generally poor fit at the semi-natural pasture site could result from these complex dynamics, particularly in comparison to the intensive pasture sites where species adjustment may be more extreme and composition more homogeneous. The semi-natural pasture sites combine more similar numbers of species from their native habitats and human adapted (largely introduced) species when compared to the intensive pasture sites, which are more weighted in terms of the latter (Table 2). The semi-natural pasture sites are also in greater proximity to forest sites, which could impact how species use these sites in terms of complex source-sink dynamics (Borges and Brown 2004; Matthews 2021). For example, perhaps some species, particularly indigenous species, use the semi-natural pasture to disperse between different fragmented forest habitats (Borges et al. 2008). The effect of combining different types or groups of species (eg. core-occasional, indigenous-introduced) into a single sample on SAD form is well known (eg. Magurran and Henderson 2003; Matthews and Whittaker 2015; Antão et al. 2017), though their effect in the context of METE remains to be explored further.

### 4.6 Summary

Across METE predictions for all species together, the forest habitats are better predicted by METE than the semi-natural pasture habitat. The intensive pasture is intermediately well fit for the SAD and MRDI, and better fit for the SAR, though the residuals are not normally distributed.

For the forest sites, the deviations from METE are comparatively small and there are less noticeable trends in the residuals. The pasture sites are characterized by a few very abundant species, which is consistent with the abundance of several small bodied introduced spiders. The semi-natural pasture is particularly poorly described by METE across metrics. This could be due to high source-sink dynamics and complex interactions between indigenous and introduced species, particularly because of the proximity to other land uses, or because of the varying levels of management in semi-natural pastures over the course of a year. The comparatively better fit at the intensive pasture site could result from the sensitive species having already been lost and thus the remaining arthropod communities comprising species that are better adapted to the high level of management intensity. In terms of METE, this may mean that state variables change less rapidly at these sites, or perhaps the more homogeneous species composition means interactions are simpler despite the higher degree of disturbance.

Analyzing the deviation from METE predictions across land use has provided us with useful information about how land use and related disturbance is affecting macroecological patterns in Azorean arthropods. We were additionally able to interpret the deviations from METE predictions ecologically. We expect this type of comparison between METE predictions and ecosystems under land management disturbance to be helpful in identifying how land use affects macroecological patterns across other habitats and taxa.

