## Supporting Information for "Land use change through the lens of macroecology: insights from Azorean arthropods and the Maximum Entropy Theory of Ecology"

#### Appendix S1 Body length to body mass conversion

To convert body length values to body mass values, we use a power equation in its arithmetic form: $M = aL^b$, where $M$ is body mass (in mg), $L$ is body length (in mm) and $a$ and $b$ are fitted constants. We source $a$ and $b$ values from empirical scaling equations in the literature for different class, orders, sub-orders, and families of arthropods (using the lowest taxonomic level we could find): Aphidae, Araneae, Blattodea, Chilopoda, Coleoptera, Dermaptera, Diplopoda, Hemiptera - Heteroptera, Hemiptera - Homoptera (non-aphid), Opilionida, Orthoptera, Psocoptera and Thysanoptera (values all taken from Hódar 1996); Pseudoscorpions (Wardhaugh 2013); Trichoptera (Baumgärtner and Rothhaupt 2003); and for all species not in those groups ($n = 3$) we use general arthropod parameter values taken from Hódar (1996). These equations only relate to adult individuals and thus we remove all juvenile individuals prior to fitting.

Note that some of the references cited above use the log–log version of the power model, whereas others (and ourselves) use the arithmetic form. Thus, in the former case we take the exponential of their provided $a$ parameter value to use in our equations. Note also that the value used for Pseudoscorpions is taken from a study of invertebrates in a tropical rainforest in Australia (and they use standard major axis regression rather than OLS); however, only two of the species in our analyses are Pseudoscorpions and thus any discrepancies due to the different focal region are unlikely to affect the results.

These scaling relationships are approximate, and using them at the level of class, order, sub-order, or family, masks likely significant variation within these taxonomic levels. Overall, the generated mass values should only be taken as very approximate estimations of the true values.

### Appendix S2 Intraspecific body mass variation

The body length measurements in our main dataset are averages and for many species the original individual measurements (from which the averages were calculated) were not stored. However, from two additional datasets, we source body length measurements for multiple individuals for a number of Coleoptera species ($n = 26$) and Araneae ($n = 26$) species that are present in our main dataset. This allows us to create body mass values for individuals and thus assess intraspecific variation in body mass, and finally convert this into a variance in metabolic rate which we use to reconstruct more realistic metabolic rate distributions.

Spider data comes from Macías-Hernández et al. (2020), and all measurements are from individuals sampled in the Azores. Body length is measured as mm. For all but two species we have measurements from males and females (median number of individuals across species = 5 males and 5 females). Body mass for each individual is calculated using the same approach described in SI Appendix S1. Note that although this data has separate body length values for female and male spiders, in the main dataset we have only average body length, and therefore we will use only the overall mean and variance for spiders without separating by sex.

Coleoptera data comes from Terzopoulou et al. (2015), and comprises measurements of numerous individuals per species (median = 10.5), all sampled in the Azores; the sex of individuals was not recorded. It is important to note that these Coleoptera measurements differ from those in the main dataset in that they were made using dried individuals (the main dataset measurements used individuals stored in ethanol). However, as all the individuals in this intraspecific analysis were dried, and we do not compare directly with the values in the main dataset, this should not be an issue. We use the length of head + pronotum + elytra, except for species in the Staphylinidae where we also include length of the abdomen (i.e., head + pronotum + elytra + abdomen).

Coleoptera and Araneae are the two most common orders in the data, and therefore it is the most important to have an idea of intraspecific variation in body mass for these orders. For less abundant species, this variation matters less as there are only a few of each species present in the data. For all other orders in the dataset, we use the results of the Coleoptera variation.

We plot the body mass distributions for the four most abundant species of Coleoptera and Araneae in this dataset in Fig. S1, overlayed with the best fit normal distributions. We find that these distributions can be roughly approximated by normal distributions. Gouws et al. (2011) also found that intraspecific body size distributions are usually approximately normal across many species of insects, including beetle species. Note that for spiders we find that this distribution is much more likely to be bimodal because of the sexual dimorphism present in many of the species. However, as mentioned above, the full dataset provides only one average body length regardless of sex, and so we approximate the spiders body mass distribution as a single normal distribution, as with the beetles. This approximation may be closer to representing the female spiders as we are more likely to sample a larger number of them due to their longer life span (Wise 1995). However, males are also well represented in the samples since pitfall traps tend to capture more active individuals, which is the case for male spiders looking for a mate.

We plot the relationship between the $\log_{10}$ of the mean and the $\log_{10}$ of the variance for both beetles and spiders in Fig. S2. The slopes, intercepts, and $R^2$ correlation coefficient values are shown in Table S1. We use these values to simulate variation in body mass for all species in our dataset and to reconstruct empirical MRDIs.

For both the beetles and the spiders, we find that the relationship between the $\log_{10}$ of the mean body mass and the $\log_{10}$ of the variance is roughly linear, without an obvious trend in the residuals. The slopes and intercepts are similar for the beetles and spiders, although the slope for the spiders is slightly steeper ($2.22 \pm 0.13$ versus $1.99 \pm 0.12$). This means the variance will be higher for spiders than for most species with comparable mass, which makes sense given that we expect the true distribution for the spiders to be bimodal.

To then reintroduce intraspecific body mass variation in the main dataset, we simulate these distributions. For each species, we draw $n_0$ samples from the corresponding normal distribution for body mass. Because the samples are drawn randomly, some samples for low mass species will be below zero, even though mass must be strictly positive. We fix this by setting these draws to the smallest positive mass plus a small amount of noise (1% of the smallest mass multiplied by a normal distribution with mean equal to zero and variance equal to one, $0.01 m_{\text{minimum}}\, \mathcal{N}(\mu = 0, \sigma^2 = 1)$) to avoid duplicate values of the smallest mass. We use the relationship for beetles for all orders except for Araneae, where we use the relationship we obtained for spiders. Some of the orders may be more similar to spiders, or be quite different overall, but since Coleoptera and Araneae are the two most common orders in the dataset, differences among other orders should not overly impact the analysis.

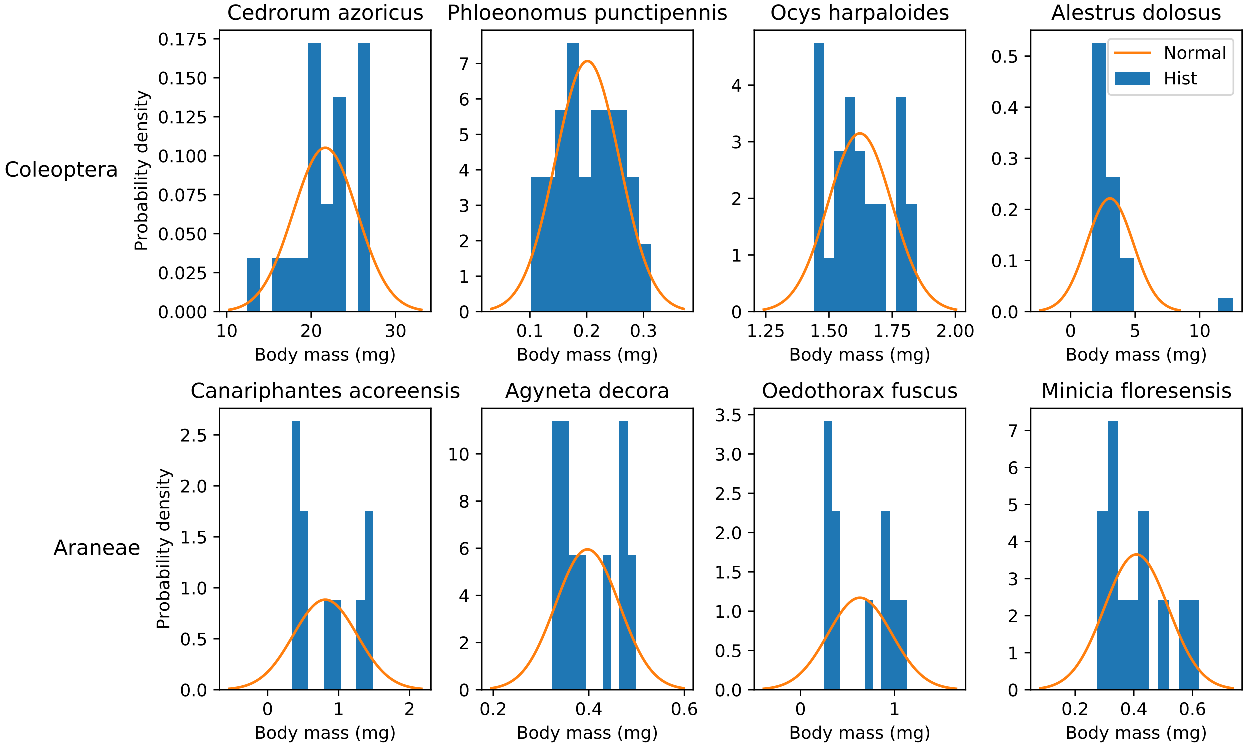


Figure S1: Histograms and best fit normal distributions for the four most abundant species of Coleoptera and Araneae present in the data that includes intraspecific variation.

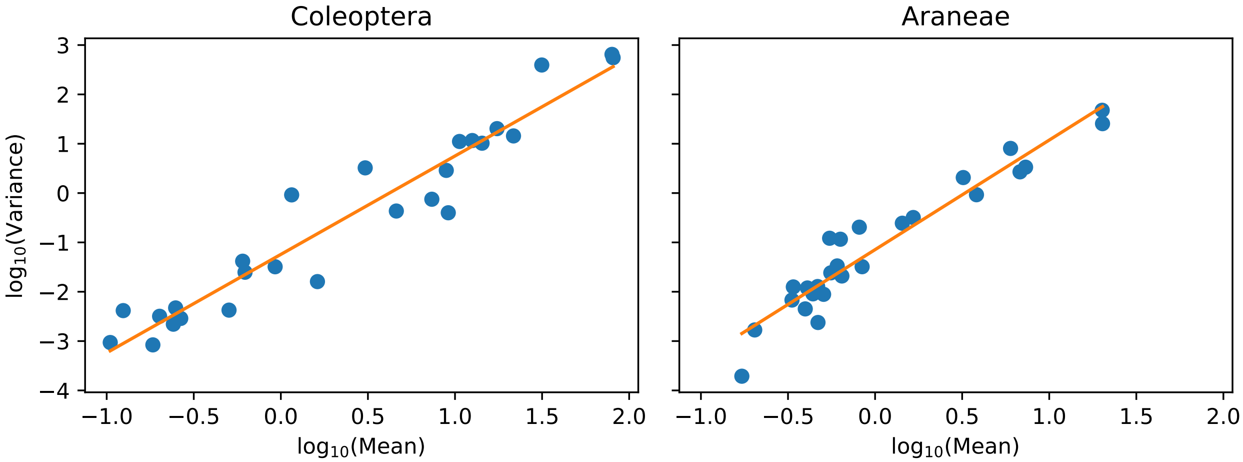


Figure S2: $\log_{10}$ of variance versus $\log_{10}$ of mean body mass for data that includes intraspecific body mass variation for both Coleoptera and Araneae.

| Order | Slope | Intercept | $R^2$ |
|---|---|---|---|
| Coleoptera (beetles) | 1.99 ± 0.12 | -1.24 | 0.925 |
| Araneae (spiders) | 2.22 ± 0.13 | -1.15 | 0.919 |

Table S1: Results from the regression of $\log_{10}$ of variance versus $\log_{10}$ of mean body mass for individuals where data is available.

### Appendix S3 METE Review

METE predicts many different macroecological patterns simultaneously by maximizing Shannon information entropy given a set of constraints (Harte et al. 2008; Harte 2011; Harte and Newman 2014; Brummer and Newman 2019). Its core distribution is the ecological structure function $R(n, \varepsilon | S_0, N_0, E_0)$, which is a joint distribution over abundance $n$ and metabolic rate $\varepsilon$ given the number of species $S_0$, the number of individuals $N_0$, and the total metabolic rate $E_0$. Thus, $R d\varepsilon$ is the probability that an individual picked at random from all species with abundance $n$ has metabolic rate between $\varepsilon$ and $\varepsilon + d\varepsilon$. Note that $n$ is discrete, and $\varepsilon$ is continuous. In practice, we scale the metabolic rate such that the smallest metabolic rate in the ecosystem has $\varepsilon = 1$.

We use the method of Lagrange multipliers to maximize the information entropy $\sum_n \int_\varepsilon d\varepsilon R \log(R)$ given the following constraints:

$$\begin{aligned} \frac{N_0}{S_0} &= \sum_{n=1}^{N_0} \int_{\varepsilon=1}^{E_0} d\varepsilon \; nR(n, \varepsilon) \\ \frac{E_0}{S_0} &= \sum_{n=1}^{N_0} \int_{\varepsilon=1}^{E_0} d\varepsilon \; n\varepsilon R(n, \varepsilon). \end{aligned} \tag{S1}$$

We additionally require the distribution $R$ to be normalized such that $\sum_{n=1}^{N_0} \int_{\varepsilon=1}^{E_0} d\varepsilon \; R(n, \varepsilon) = 1$. The solution for the ecological structure function is

$$R(n, \varepsilon | S_0, N_0, E_0) = \frac{\exp\left(-\lambda_1 n - \lambda_2 n \varepsilon\right)}{Z}, \tag{S2}$$

where the Lagrange multipliers $\lambda_1$ and $\lambda_2$ are solved from the constraints, and the normalization $Z$ is calculated as $\sum_{n=1}^{N_0} \int_{\varepsilon=1}^{E_0} d\varepsilon \; \exp\left(-\lambda_1 n - \lambda_2 n \varepsilon\right)$. To a very good approximation given typical empirical values for $S_0$, $N_0$, and $E_0$ (Harte 2011; Brummer and Newman 2019), $\lambda_2 = S_0/(E_0 - N_0)$, and $\lambda_1$ can then be solved from

$$\frac{N_0}{S_0} = \frac{\sum_{n=1}^{N_0} e^{-\beta n}}{\sum_{n=1}^{N_0} e^{-\beta n}/n}, \tag{S3}$$

where $\beta = \lambda_1 + \lambda_2$. The distribution $R$ can then be used to derive other macroecological distributions.

#### Appendix S3.1 Species abundance distribution

We obtain the METE SAD prediction $\Phi(n)$ by integrating the structure function over $\varepsilon$. The METE prediction is equivalent to the max likelihood prediction for the log series (White et al. 2012, Appendix A). This prediction assumes that the number of species is large enough that we can ignore certain terms, which eliminates any dependence on $E_0$. The resulting prediction is the log series distribution

$$\Phi(n | S_0, N_0) = \frac{e^{-\beta n}}{n \log\left(1/\left(1 - e^{-\beta}\right)\right)} \tag{S4}$$

#### Appendix S3.2 Metabolic rate distribution of individuals

The METE MRDI prediction $\Psi(\varepsilon)$ is obtained by summing the structure function multiplied by $n$ over $n$ and correcting the normalization,

$$\Psi(\varepsilon) = \frac{S_0}{N_0} \sum_{n=1}^{N_0} nR(n, \varepsilon). \tag{S5}$$

This sum gives

$$\Psi(\varepsilon | S_0, N_0, E_0) = \lambda_2 \left(e^{\beta} - 1\right) \frac{e^{-\gamma}}{(1 - e^{-\gamma})^2}, \tag{S6}$$

where $\gamma = \lambda_0 + \lambda_1 \varepsilon$. Note that we use a slightly different form for $\Psi$ compared to Eq. 7.33 in Harte (2011), where $\beta$ has been replaced with $e^{\beta} - 1$. The normalization of $\Psi(\varepsilon)$ in Eq. S6 is significantly better, as

$\int_\varepsilon d\varepsilon\ \Psi(\varepsilon)$ is much closer to 1. This form still allows the cumulative distribution function and the rank ordered distribution to be solved analytically, and is numerically very similar to the full expression without any approximations, even for analyses at the transect level.

#### Appendix S3.3 Species–area relationship

The SAR can be predicted by combining the SAD with the species-level spatial abundance distribution (SSAD) $\Pi(n|A, A_0, n_0)$, which predicts the number of individuals present in an area $A$ given $n_0$ individuals of that species in a larger area $A_0$. The number of species at a given scale $A$ can be predicted by multiplying the SAD by the probability that a species with abundance $n_0$ is present at that scale and then summing over $n_0$,

$$S(A) = \sum_{n_0=1}^{N_0} \Phi(n_0)\left(1 - \Pi(0|A, A_0, n_0)\right). \tag{S7}$$

The SSAD can also be predicted by maximizing information entropy given the constraint $\sum_{n=0}^{n_0} n\Pi(n) = n_0 A/A_0$. The solution corresponds to the finite negative binomial distribution (Conlisk et al. 2007; Zillio and He 2010)

$$\Pi(n|A, A_0, n_0) = \frac{\binom{n+k-1}{n}\binom{n_0-n+kA_0/A-k-1}{n_0-n}}{\binom{n_0+kA_0/A-1}{n_0}}, \tag{S8}$$

with aggregation parameter $k = 1$ (Harte 2011; Wilber et al. 2015).

METE also predicts that all nested SARs will collapse onto a single universal curve when plotted as the slope of the SAR $z$ versus $D = \log(N_0/S_0)$, a scale parameter (Harte 2011; Wilber et al. 2015).

### Appendix S4 Broader community level analysis

We feel that our analysis treating the transects as replicates across land use is stronger, particularly when comparing data to METE. This is because METE predictions are made within a single community and aggregating data over disparate locations, even of the same land use, may create a mismatch between the theory expectations and the aggregated data. Additionally, the number of species scales differently when summing multiple small patches than in a large patch of comparable area.

Despite that, we present the analysis at the community level here. In this case, all transects with the same land use are aggregated together, and we compare that empirical data to the METE prediction made with the total number of species and individuals for that land use. The mean least squared error for the SAD and MRDI across land uses is shown in Fig. S3. These results are similar to those obtained when the transects are analyzed individually, though note here that the MRDI is the worse fit at the intensive pasture rather than the semi-natural pasture. Another difference is that the MRDI is comparatively better fit than the SAD at the forest sites. This is primarily as the SAD is significantly worse fit at the community level. The semi-natural pasture is still the only site that is poorly fit by both metrics. This fits with our interpretation in the main text that this site is the most poorly described by METE. The SAD results in particularly are very similar when analyzed at the community level.

We also show the empirical rank ordered SADs along with the corresponding METE predictions in Fig. S4. Note that again the pasture sites are characterized by a few very abundant species, and METE under predicts the number of singletons across sites.

The empirical rank ordered MRDIs along with the corresponding METE predictions in Fig. S5. Again here, METE over predicts the metabolic rate of the highest metabolic rate individuals. This is particularly unsurprising here, as METE will predict higher metabolic rate individuals in larger ecosystems, which we have created here by aggregating across transects. It is likely that the maximum size of arthropods is more constrained at the transect level than at this larger community level.

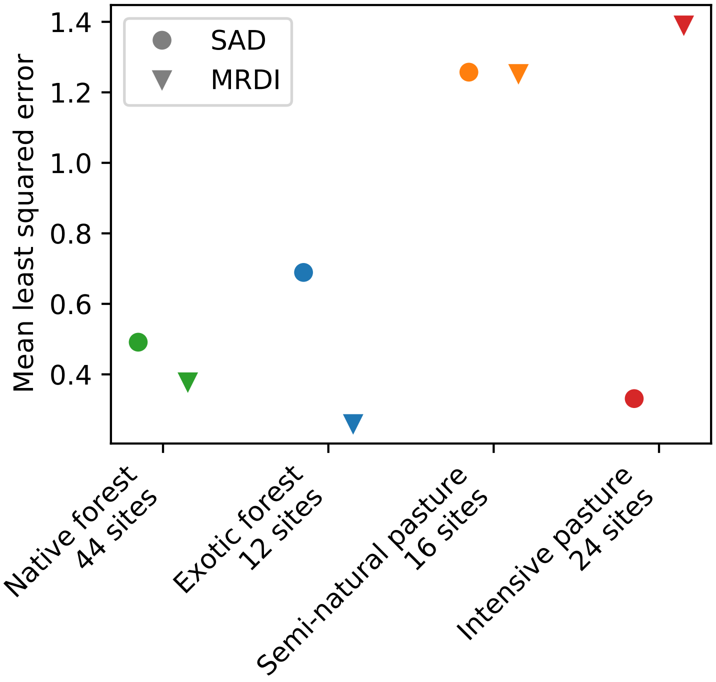


Figure S3: The mean of the the mean least squared error for the SAD and the MRDI across land uses when transects are aggregated rather than analyzed individually. There are no error bars as there are no replicates when the data are analyzed this way.

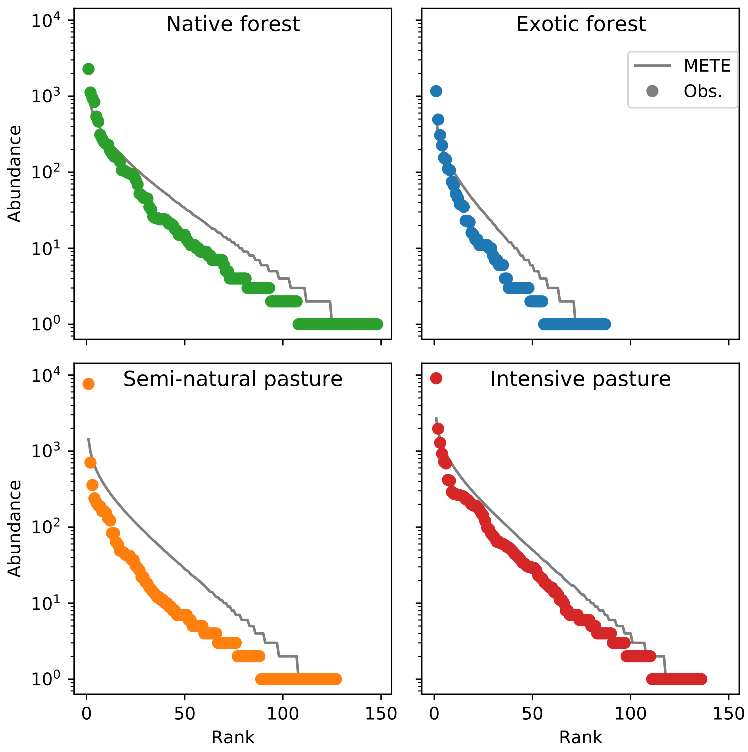


Figure S4: Aggregated rank ordered SADs by land use. The solid line is the METE prediction and the points are observed rank abundance.

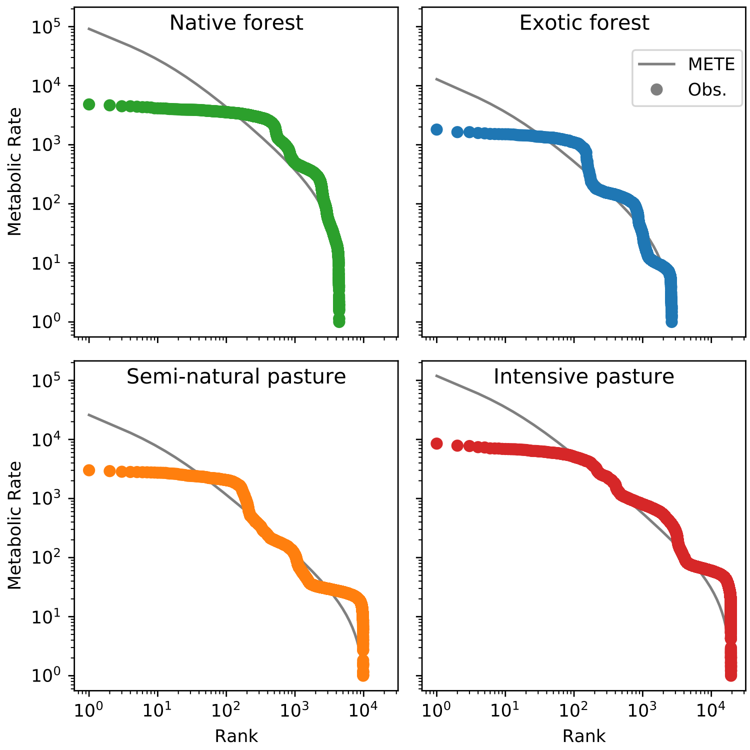


Figure S5: Aggregated rank ordered MRDIs compared to METE predictions by land use. The empirical curve here is simulated from assuming a mean-variance relationship and adding variance to the mean body mass for each species.

#### Appendix S5 Kolmogorov-Smirnov test

As a second goodness of fit test to check our results using mean least squares, we use the Kolmogorov-Smirnov test comparing the empirical CDF and the METE predicted CDF. For the SAD, the KS test must be adapted as the distribution is discrete with many duplicate values (ie. many singletons). We made use of the R package DGOF, which implements the KS test for discrete distributions (Arnold and Emerson 2011). This is not an issue for the MRDI as it does not have repeated values. We plot the mean and standard error of the test statistic $D_{KS}$ for both the SAD and MRDI for each land use in Fig. S6.

The results here are comparable to those obtained with mean least squares. The MRDI is worse fit than the SAD, and the semi-intensive pasture is the worst fit for both the SAD and the MRDI. The intensive pasture results are also similar as it is among the best fit for the SAD, and intermediately well fit for the MRDI. That these results are comparable to the mean least squares results is important, as our conclusions about how well METE describes the data across land use do not appear to depend on the goodness of fit test itself.

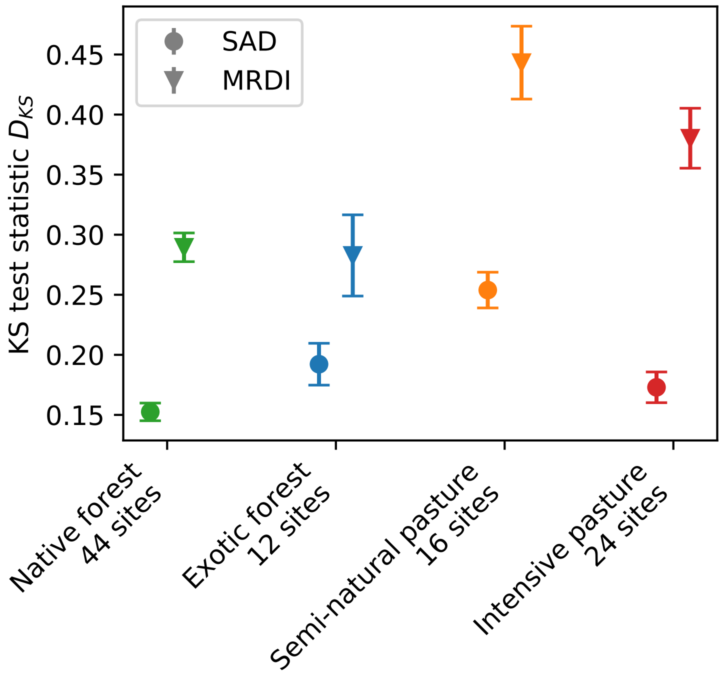


Figure S6: The mean and standard error of the KS test statistic $D_{KS}$ for the SAD and MRDI across transects for each land use.

#### Appendix S6 SAR comparison of number of species

In addition to the comparison of slopes in the main text, we can compare the predicted number of species at each scale directly. The mean least squares across transects at each land use are shown in Fig. S7, again as the mean at the land use with its standard error. The direct plots of $\log(S)$ versus $\log(A)$ do not show the difference between the theoretical predictions and the observed values very well as the differences are relatively small, so we instead show the residuals of $\log(S_0)$ for each transect in Fig. S8. Note that as mentioned in Methods, the largest scale corresponds exactly in all cases, and so we really only have 7 points of prediction in this case. Here again we see that METE over predicts $S_0$ at smaller scales for the pasture sites, and the residuals are relatively randomly distributed for the forest sites.

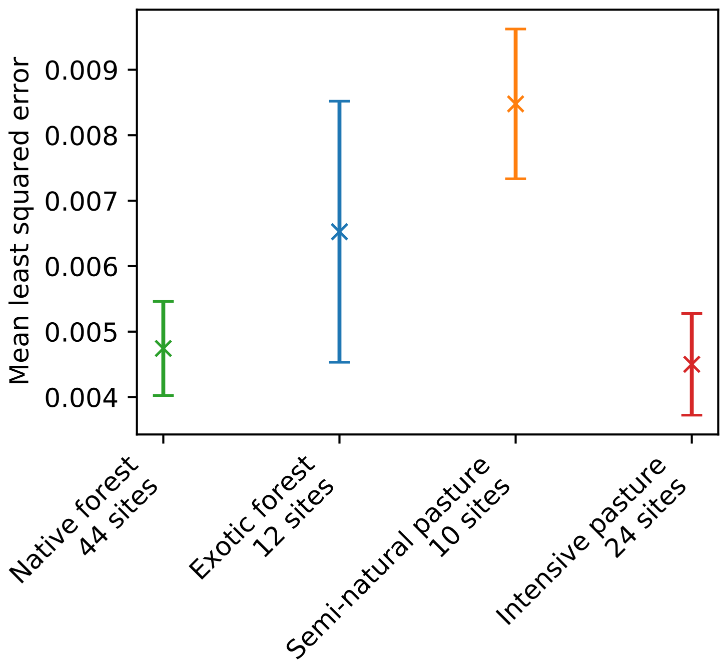


Figure S7: The mean and its standard error for the mean least squares of the predicted number of species for each transect, organized by land use.

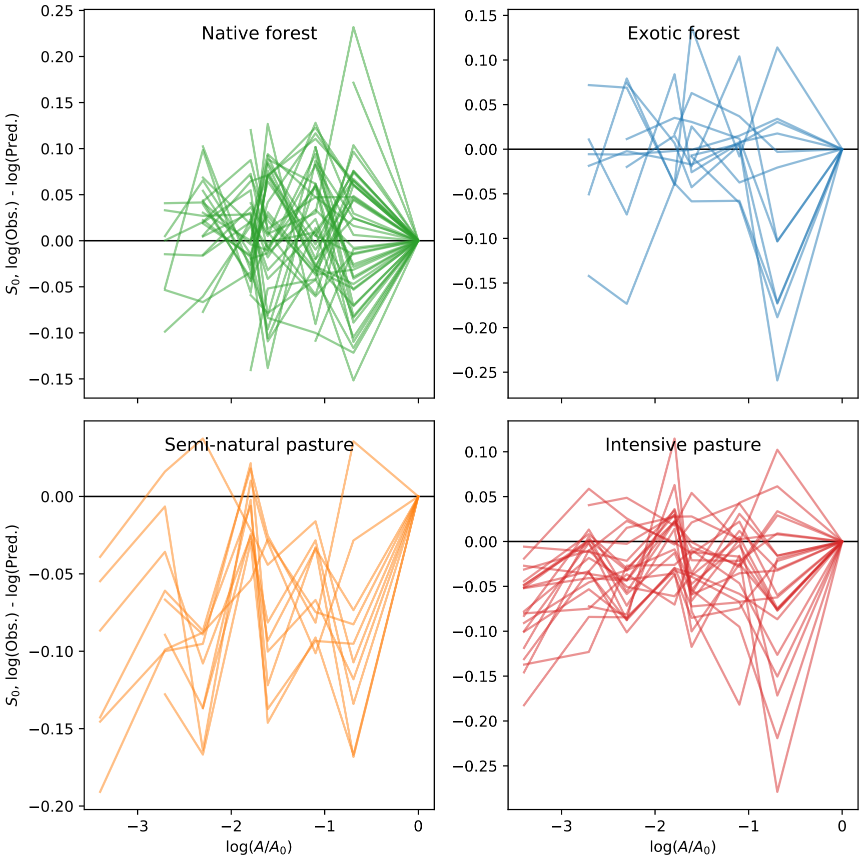


Figure S8: The observed minus the predicted $\log(S_0)$ at each scale $\log(A/A_0)$. Each line traces out a single transect.

### Appendix S7 Additional information on comparing METE predictions to data

#### Appendix S7.1 Species abundance distribution (SAD)

There are many existing methods for comparing empirical SADs to data; however, there is not a single optimum goodness of fit metric (Connolly and Dornelas 2011; Matthews and Whittaker 2014). In general, binning should be avoided as the shape of the distribution depends on the binning interval, and given the sparse data each observed bin is unlikely to have a sufficiently large expectation to have a meaningful $\chi^2$ test (Williamson and Gaston 2005; Gray et al. 2006; Ulrich et al. 2010). Log-likelihood based methods are also inappropriate in this case as we are not trying to determine a preferred model and instead are looking for a goodness of fit test.

Given this, we use the mean least squares of the rank ordered natural log of abundance as our primary goodness of fit metric to compare the METE predictions to data. Mathematically, this means we take $\sum_{i=1}^{S_0} \left(\log\left(n_{i,\text{observed}}\right) - \log\left(n_{i,\text{predicted}}\right)\right)^2 / S_0$, where $n_i$ is the abundance of the species with rank $i$, and we take the mean over all $S_0$ ranks. Although Matthews and Whittaker (2014) note that mean least squares of rank ordered data violates some underlying statistical assumptions, namely that the data points are not statistically independent (see also Connolly and Dornelas 2011), we still find it preferable to their recommended method of using a parametric bootstrap. For many test statistics this method scales with the number of points, making comparisons between land use types challenging. As that is our primary objective here, we prefer to use the mean least squares.

To ensure our results are robust to our choice of goodness of fit metric, we additionally performed Kolmogorov-Smirnov (KS) tests for each transect and obtained the two-sided test statistic $D_{KS}$ when the empirical cumulative distribution function (CDF) is compared to the METE predicted CDF. These results can be found in Appendix S5, and are very similar to the results obtained by mean least squares. For the KS test for the SAD, we used the R package DGOF which implements the KS test for discrete distributions (Arnold and Emerson 2011).

#### Appendix S7.2 Metabolic rate distribution of individuals (MRDI)

We first reintroduce intraspecific body mass variation as described in Appendix S2. For adult arthropods, we then use metabolic scaling to convert body mass data to metabolic rate. We assume that $\varepsilon \propto m^{3/4}$, where $m$ is the body mass. The metabolic rates are then scaled such that the smallest $\varepsilon = 1$. We then rank order the data in order to compare to the METE prediction.

As with the SAD, we primarily use the mean least squares of the rank ordered natural log of metabolic rates as a goodness of fit metric for each land use. In this case, that means $\sum_{i=1}^{N_0} \left(\log\left(\varepsilon_{i,\text{observed}}\right) - \log\left(\varepsilon_{i,\text{predicted}}\right)\right)^2 / N_0$, where $\varepsilon_i$ is the metabolic rate of the individual with rank $i$ and the mean is now over the number of individuals $N_0$. Alternative options for this comparison would be $R^2$ as defined in eg. Xiao et al. (2015), or to bin the data and use a $\chi^2$ test. However, binning relies on a large number of points per bin that are not available here at the individual transect level, and the results can depend on the bin width. In order to test the robustness of our goodness of fit metric we additionally perform Kolmogorov-Smirnov goodness of fit tests for the empirical CDF compared to the METE predicted CDF. Again, we obtain similar results to the mean least squared analysis (Appendix S5).

#### Appendix S7.3 Species–area relationship (SAR)

Along each transect, there are 30 individual pitfall traps arranged linearly. For each transect individually, we compare the resulting empirical SAR to the METE prediction by first averaging the number of species at different scales. We choose scales of 1, 2, 3, 5, 6, 10, 15, and 30 traps. These relative scales were chosen as they use all of the data available at every scale, or put another way, these numbers are all factors of 30. This is slightly different than has been done in other comparisons, which use a number of cells that is a factor of two and divide repeatedly in half (eg. Franzman et al. 2021).

To compare these average numbers of species to the METE prediction, we use the functions for the log series SAD $\phi(n)$ and the finite negative binomial SSAD $\Pi(n)$ from the `macroeco` software package (Kitzes et al. 2015; Kitzes and Wilber 2016), which we have adapted for use with Python 3.0.

There are many different methods for comparing the predicted SAR to data. We could fix the number of species and individuals at the largest scale and predict the number of species at every smaller scale from this anchor scale, or we could compare the predicted slope $z$ of the SAR against the scale parameter $D = \log(N_0/S_0)$, which as noted above collapses SARs onto a single, universal curve (Harte et al. 2009; Wilber et al. 2015). Harte (2011) provides an equation for $z$ given the empirical number of species and individuals at a given scale, assuming that we bisect the plot in two. We use a similar approach, in that we use the number of species and individuals at a given scale to predict the number of species at the next smallest scale in consideration and then use that to predict a slope. Mathematically, we write

$$z_i = \frac{\log(S_i/S_{i-1})}{\log(A_i/A_{i-1})}, \tag{S9}$$

where $i$ indexes the scale. In the list above then, $i = 1$ corresponds to the average number of species at the scale of 1 cell, and $i = 8$ corresponds to the number of species in all 30 cells.

Given we are predicting the number of species directly using this approach, we can also compare the SAR directly (see Appendix S6). However, given the collapse onto a single curve in the $z - D$ plots we prefer that comparison. We obtain the empirical slope by comparing the number of species at the scale being considered to the number of species at the next smallest scale. This is similar to the theoretical prediction, but now the number of species at the smaller scale is also empirical. This method allows us to compare slopes at all except the smallest scale, as we do not have a smaller scale with which to make the empirical slope prediction.

We therefore have seven data points to make the comparison and again use the mean least squares between the METE prediction and the empirical data, here as $\sum_{i=2}^{8} \left(z_{i,\mathrm{observed}} - z_{i,\mathrm{predicted}}\right)^2/7$, where $i$ indexes the scale and we do not compare at the smallest scale $i = 1$. Note that for many transects we will have fewer than seven data points as we additionally only use scales where the empirical average for the number of species is greater than four ($S_0 > 4$). This is because several METE simplifications break down for small $S_0$, including the fact that we can ignore $E_0$ and derive $\beta$ from only $N_0$ and $S_0$. This means that transects with lower abundance will have fewer than seven points of comparison.

### Appendix S8 Significance testing

To support our interpretation of Fig. 1a, we perform statistical analyses showing our results are significant. The goal here is then to determine which land uses produce statistical significant differences for each pattern studied. We show that our results are robust across two statistical tests that make different assumptions about the underlying distributions, and that these results are consistent with our interpretation in the main text. We want to emphasize that given the exploratory nature of this study, these statistical analyses should not be interpreted as rigorous hypothesis testing, but instead as supporting evidence for the discussion in the main text.

First, we use is the ANOVA $F$-test and the post-hoc Tukey test to see which means are significantly different across land use for each pattern in Fig. 1a. The distribution for the mean least squared errors across transects do not appear normally distributed for any of the three patterns, which we tested using the Shapiro test, nor do they have equal variance across land uses, which we tested using the Levene test. As these are required assumptions for the $F$-test, we therefore take the log transform of the mean least squared error distributions for each pattern in Fig. 1a. We find that these distributions are now consistent with the normal distribution according to the Shapiro test and that the variances at each land use are not distinguishable according to the Levene test.

We perform the ANOVA $F$-test and find that the means of the mean least squared error are significantly different for both the SAD ($p = 10^{-8}$) and the MRDI ($p = 3 \times 10^{-4}$), but not the SAR ($p = 0.14$). We then use the Tukey test to determine which land uses have significantly different means. For the SAD, we find that the error at the semi-natural pasture is significantly different from that at all other sites, and that the error at the intensive pasture is significantly different to the error at the native forest. In other words, only the means between the exotic forest and the intensive pasture, and the exotic forest and the native forest are not significantly different. See Table S2 for the corresponding $p$-values. For the MRDI, we find that the error at both pasture sites is significantly different to that at both forest sites, but that the forest sites and the pasture sites themselves are not distinguishable from each other. See Table S3 for the $p$-values.

We note that while we have log-transformed the data before performing these tests in order to better satisfy the assumptions for the ANOVA $F$-test, the results are very similar when performed directly on the mean least squared error. In that case, the SAD and MRDI again have significantly different means across land uses. The Tukey test results for which means are significantly different are identical for the SAD, and for the MRDI the only difference is that the error at the intensive pasture is no longer significantly different from that at the forest sites; however Table S3 shows that this $p$-value was marginal at 0.047.

Finally, to show our results are robust across statistical tests, we additionally performed the non-parametric Kruskal-Wallis $H$-test to look for differences in the medians of the mean least squared error distributions, rather than the means. In this case, the only assumption is that the mean least squared errors should follow the same distribution across land uses, which should be reasonable. The results are again very similar: the SAD mean least squared errors are statistically different ($p = 5 \times 10^{-7}$), as are the errors for the MRDI ($p = 6 \times 10^{-4}$), but the median errors for the SAR are indistinguishable ($p = 0.14$).

We then use the post-hoc Dunn test to find which medians are significantly different for both the SAD (Table S4) and the MRDI (Table S5). The results for the SAD are the same again in that the median error at the semi-natural pasture is different to all other sites, and the median error at the intensive pasture and the native forest are different. For the MRDI, the median error at the semi-natural pasture is again different to both forest sites, and the median error at the intensive pasture is now different to the native forest but not the exotic forest.

Overall, these results are robust across statistical tests and are completely consistent with our interpretation in the main text that the semi-natural pasture is the worst fit across patterns, and that the pasture sites in general are worse fit than the forest sites. It also supports our conclusion that the intensive pasture is somewhat intermediate, in that for the SAD it is distinguishable from the semi-natural pasture and the native forest but not from the exotic forest, and for the MRDI it is distinguishable from both forest sites for the Tukey test on the log transformed data, or just from the native forest for the Dunn test. Additionally, while the mean least squared errors for the SAR are not statistically significantly different across land uses in any test, we can still see a clear bias of over prediction at larger scales in Fig. 4. This difference may not appear here as the error bars are quite large, and because the mean least squared error is only sensitive to the absolute deviation from the predicted pattern.

| Tukey test, SAD | Native forest | Exotic forest | Semi-natural pasture | Intensive pasture |
|---|---|---|---|---|
| Native forest | – | 0.14 | **0.001** | **0.01** |
| Exotic forest | 0.14 | – | **0.004** | 0.9 |
| Semi-natural pasture | **0.001** | **0.004** | – | **0.001** |
| Intensive pasture | **0.01** | 0.9 | **0.001** | – |

Table S2: Results of the Tukey test applied to the means of the log transformed mean least squared errors for the SAD. Each element is the $p$-value corresponding to the difference between the means, with results with $p < 0.05$ marked in bold as significant.

| Tukey test, MRDI | Native forest | Exotic forest | Semi-natural pasture | Intensive pasture |
|---|---|---|---|---|
| Native forest | – | 0.9 | **0.002** | **0.017** |
| Exotic forest | 0.9 | – | **0.009** | **0.047** |
| Semi-natural pasture | **0.002** | **0.009** | – | 0.77 |
| Intensive pasture | **0.017** | **0.047** | 0.77 | – |

Table S3: Results of the Tukey test applied to the means of the log transformed mean least squared errors for the MRDI. Each element is the $p$-value corresponding to the difference between the means, with results with $p < 0.05$ marked in bold as significant.

| Dunn test, SAD | Native forest | Exotic forest | Semi-natural pasture | Intensive pasture |
|---|---|---|---|---|
| Native forest | – | 0.41 | $\mathbf{2 \times 10^{-7}}$ | **0.032** |
| Exotic forest | 0.41 | – | **0.043** | 1.0 |
| Semi-natural pasture | $\mathbf{2 \times 10^{-7}}$ | **0.043** | – | **0.028** |
| Intensive pasture | **0.032** | 1.0 | **0.028** | – |

Table S4: Results of the Dunn test applied to the medians of the mean least squared errors for the SAD. Each element is the $p$-value corresponding to the difference between the means, with results with $p < 0.05$ marked in bold as significant.

| Dunn test, MRDI | Native forest | Exotic forest | Semi-natural pasture | Intensive pasture |
|---|---|---|---|---|
| Native forest | – | 1.0 | **0.006** | **0.049** |
| Exotic forest | 1.0 | – | **0.013** | 0.074 |
| Semi-natural pasture | **0.006** | **0.013** | – | 1.0 |
| Intensive pasture | **0.049** | 0.074 | 1.0 | – |

Table S5: Results of the Dunn test applied to the medians of the mean least squared errors for the MRDI. Each element is the $p$-value corresponding to the difference between the means, with results with $p < 0.05$ marked in bold as significant.

#### Appendix S9 Null expectations for the residuals

The residuals in Fig. 2 and Fig. 3 can be hard to interpret, and so here we plot the corresponding null expectations for the SAD, Fig. S9, and the MRDI, Fig. S10. These are calculated by simulating the abundance or metabolic rate data for each transect. The simulations are done by pulling a random variate of the SAD or MRDI for each transect assuming that the state variables are the same as in the data and that the SAD or MRDI follow exactly the METE prediction (see Table 1). The simulated data is then rank ordered and compared to the METE prediction in the same way as the empirical data. Even with this simulated data, we expect to see scatter about the zero line as each transect is a random realization of the METE prediction. As the species richness or abundance increase, we expect this scatter to decrease. The scatter for the null expectation for both the SAD and MRDI is evenly distributed across the zero line, which is not the case with the empirical data. We can see that the patterns discussed in the main text, including the consistent under prediction of the most abundant species in the SAD at the pasture sites, and the long lines of constant slope in the MRDI, do not appear in these null predictions. These patterns in the residuals can therefore be interpreted as deviations from the METE predictions that contain information about the ecosystem.

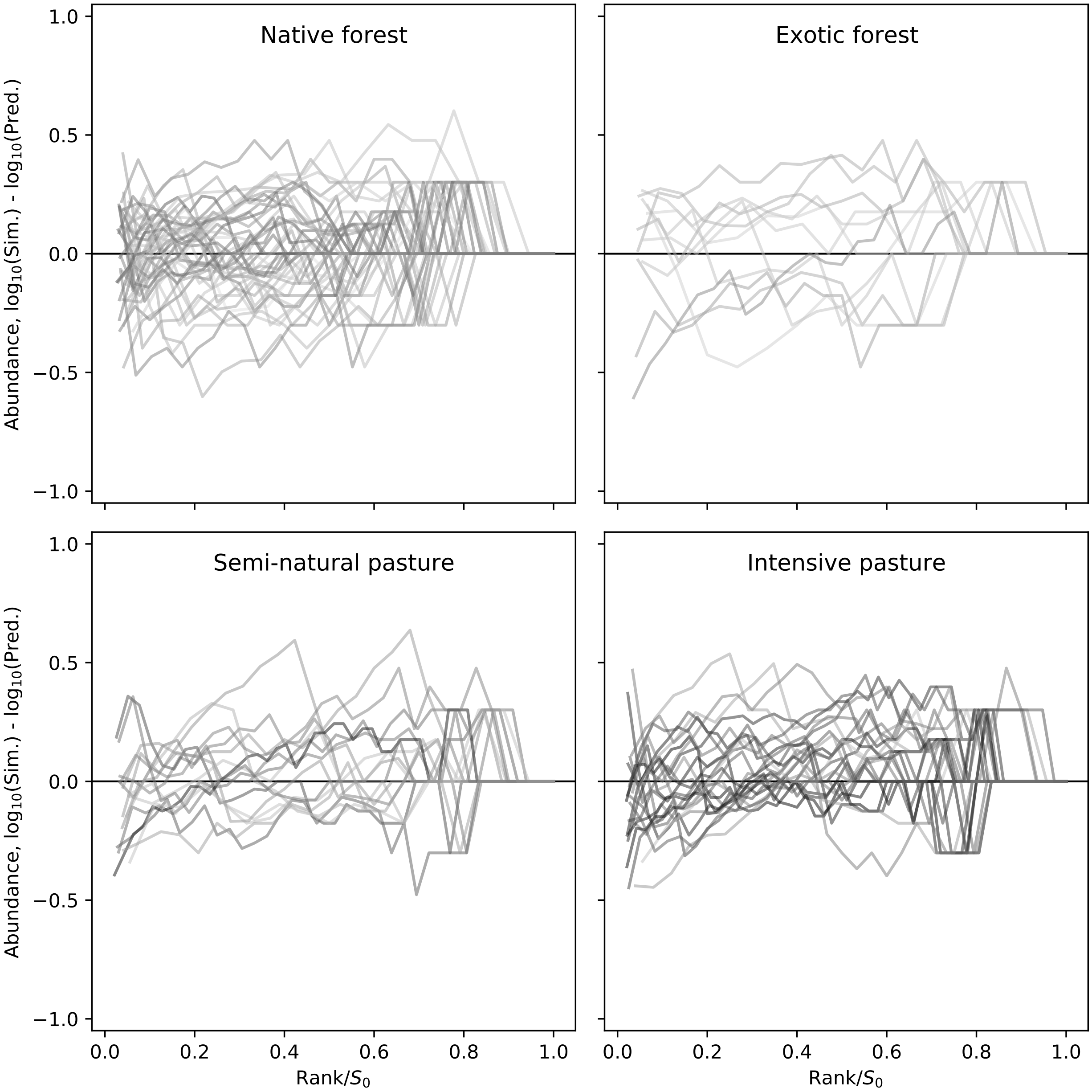


Figure S9: The residuals of simulated data for the SAD, calculated as $\log_{10}$ of the simulated abundance minus $\log_{10}$ of the predicted abundance from METE for each transect across land uses. The data is simulated by taking a random variate of the METE predicted distribution for each transect, assuming the same state variables. As the number of ranks is equal to the number of species $S_0$, the ranks on the x-axis have been rescaled by $1/S_0$ to facilitate comparison between sites. The darker lines are sites with a higher number of species, and lighter lines represent sites with fewer species.

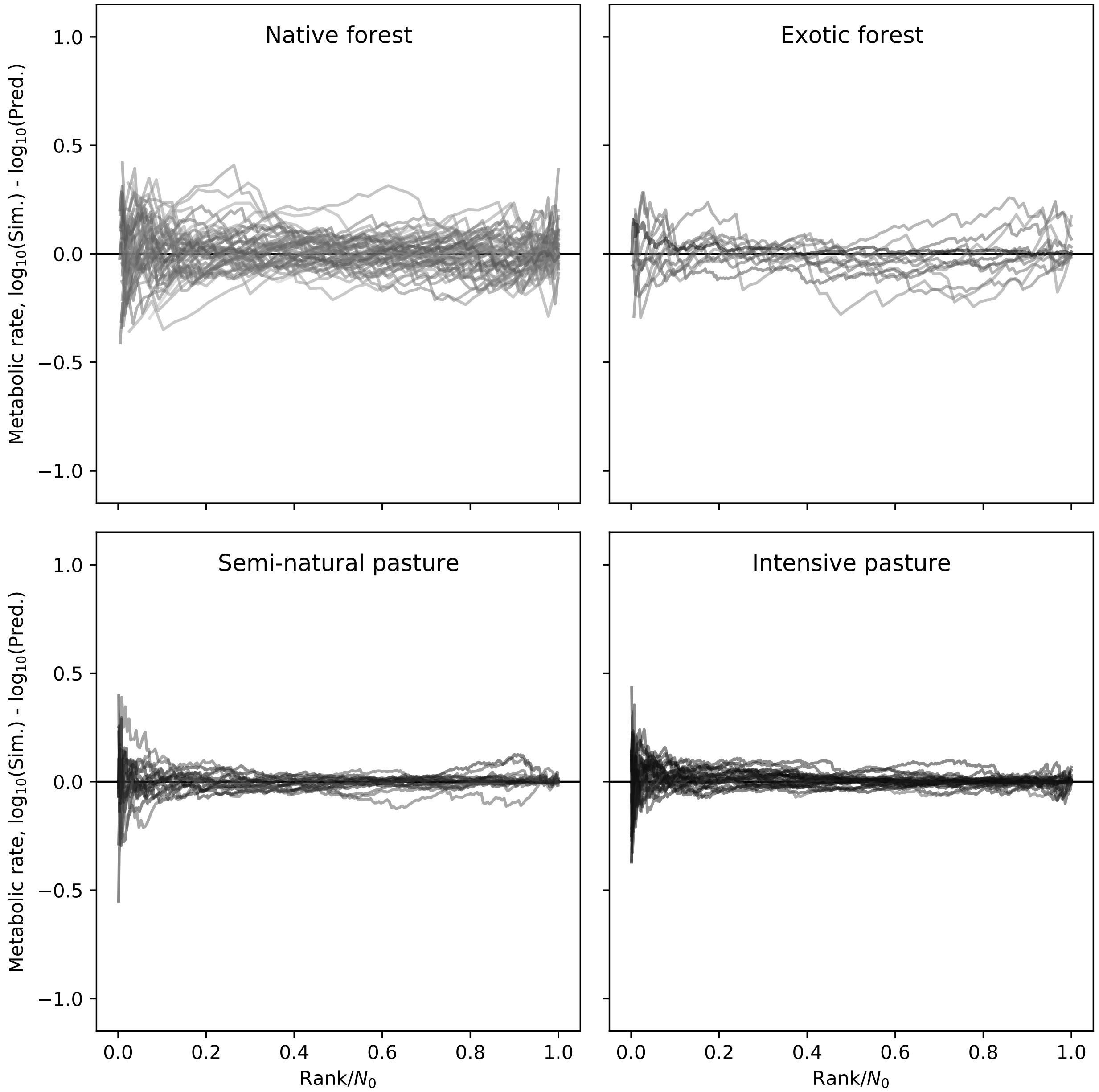


Figure S10: The residuals of simulated data for the MRDI, calculated as $\log_{10}$ of the simulated metabolic rate minus $\log_{10}$ of the predicted metabolic rate from METE for each transect across land uses. The data is simulated by taking a random variate of the METE predicted distribution for each transect, assuming the same state variables. As the number of ranks is equal to the number of species $N_0$, the ranks on the x-axis have been rescaled by $1/N_0$ to facilitate comparison between sites. The darker lines are sites with a higher number of species, and lighter lines represent sites with fewer species.

#### Appendix S10 SADs at each transect

Presented in order of increasing land use intensity:

- Fig. S11 – Native forest
- Fig. S12 – Exotic forest
- Fig. S13 – Semi-natural pasture
- Fig. S14 – Intensive pasture.

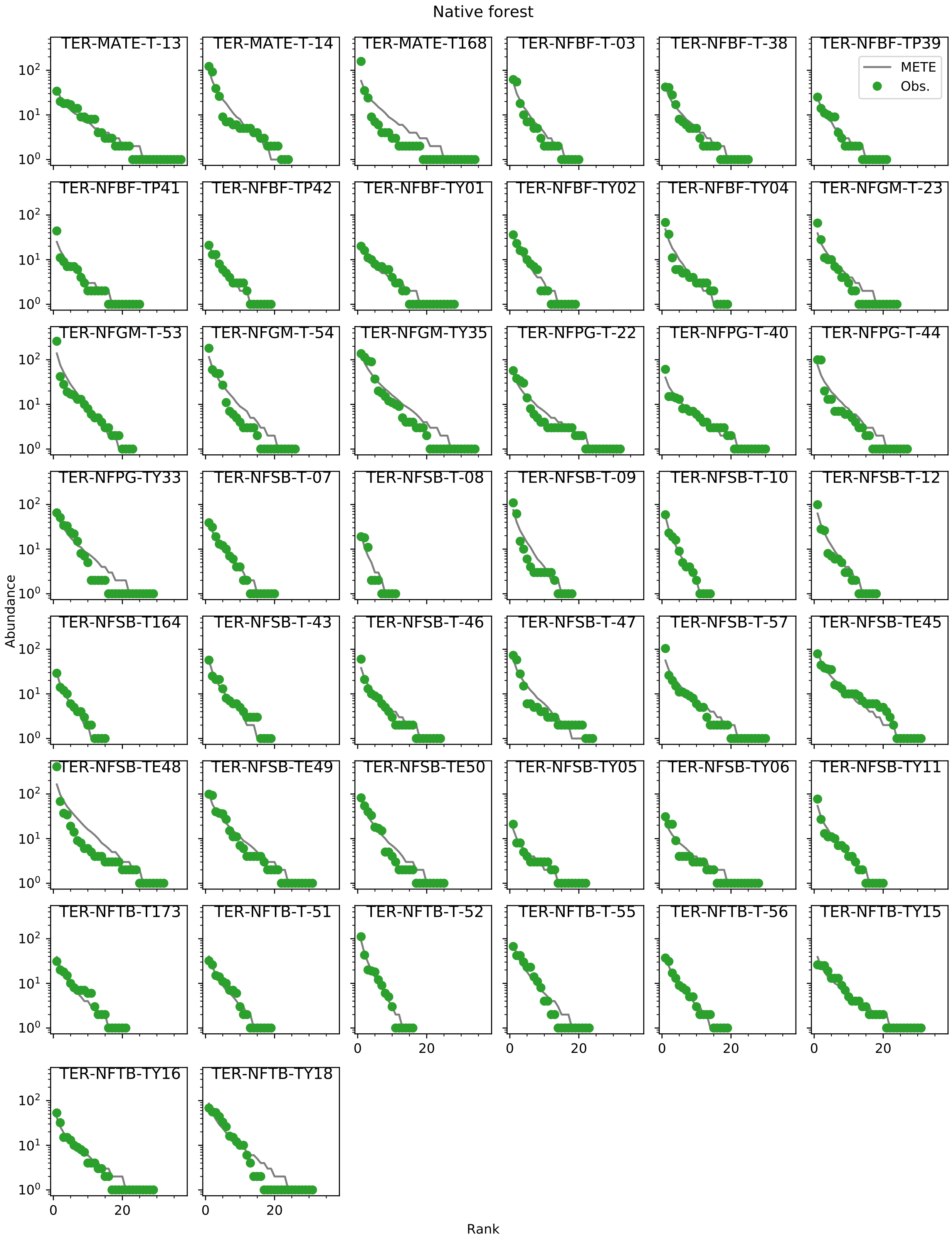


Figure S11: The rank ordered SAD at each transect in the native forest.

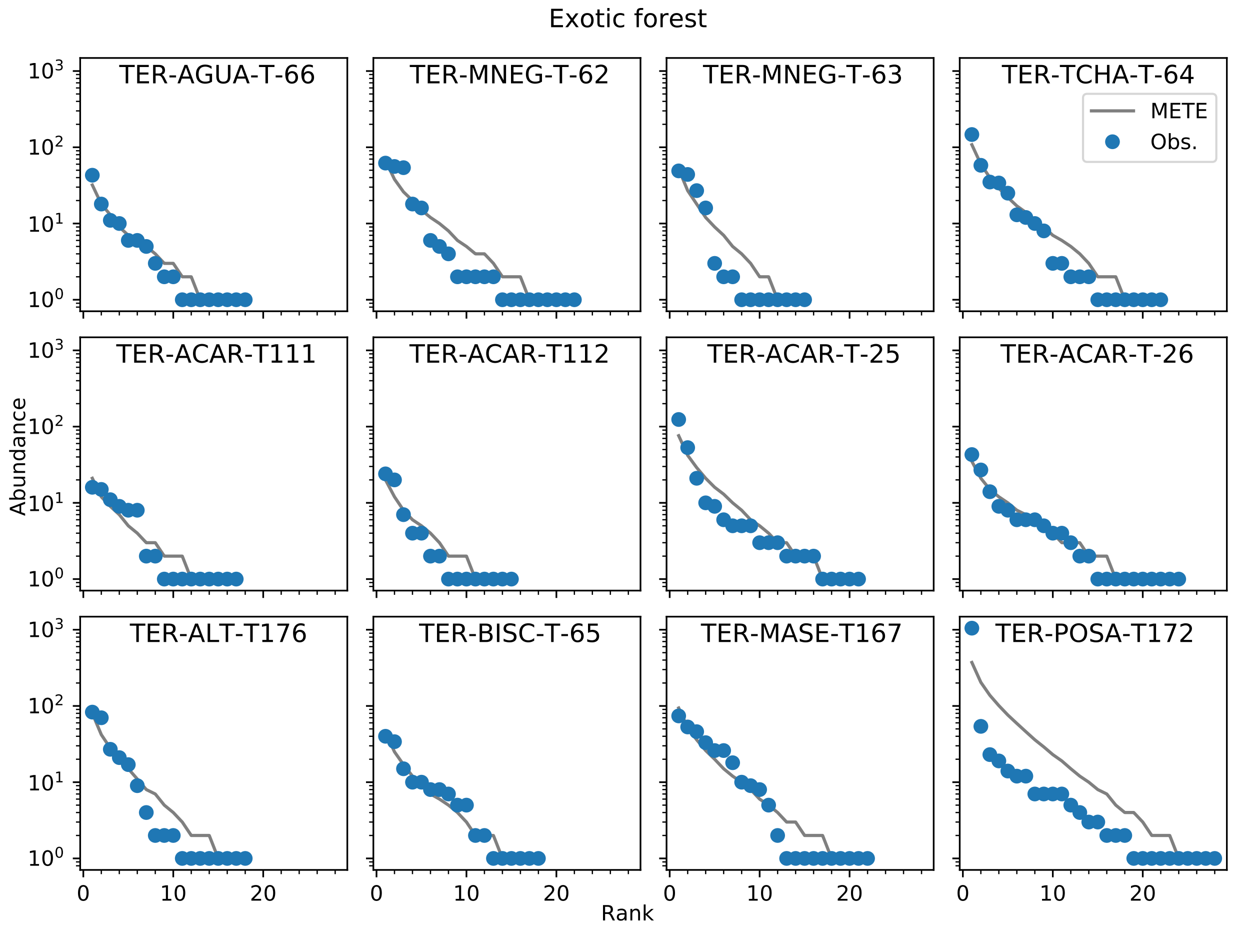


Figure S12: The rank ordered SAD at each transect in the exotic forest.

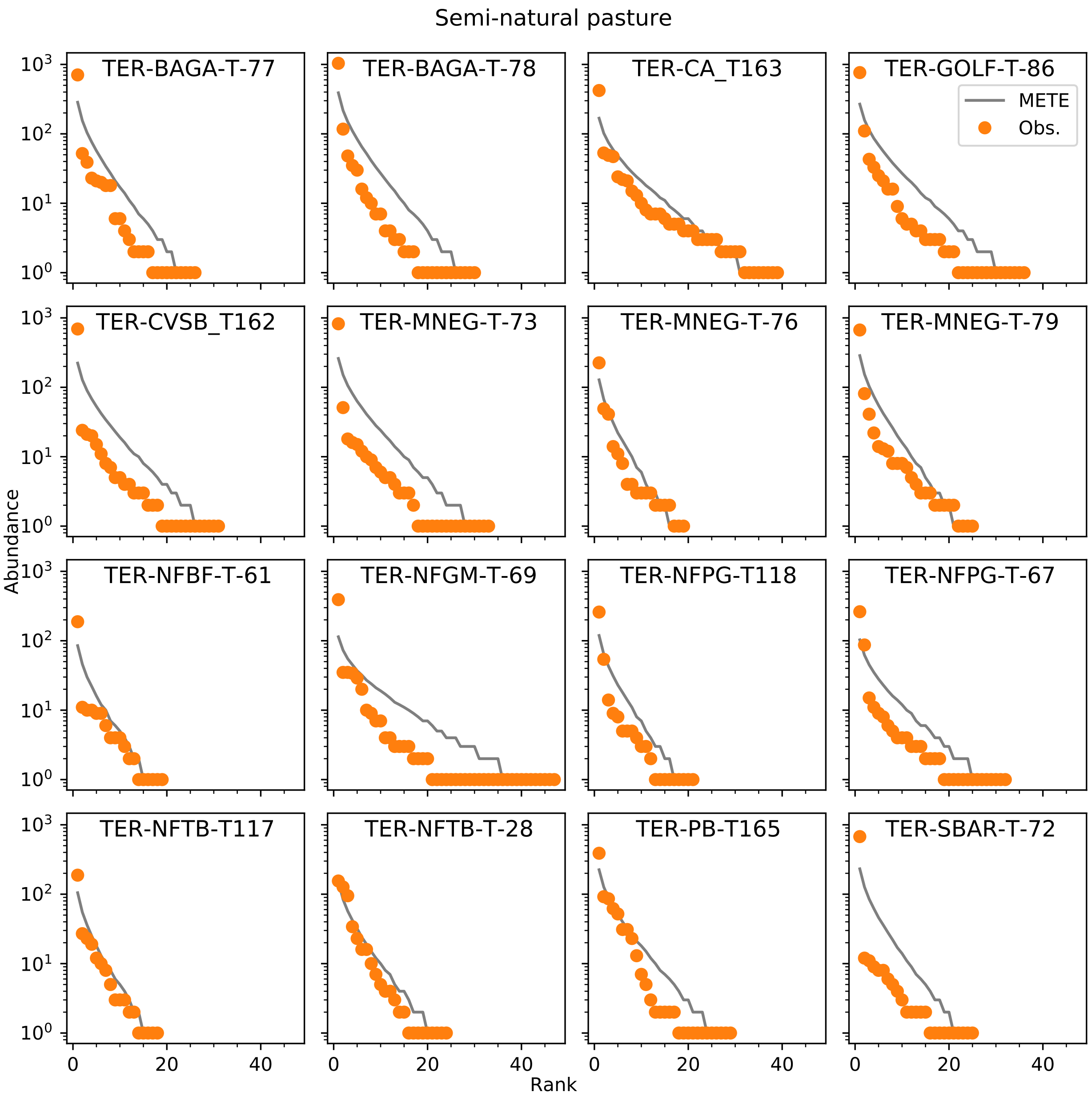


Figure S13: The rank ordered SAD at each transect in the semi-natural pasture.

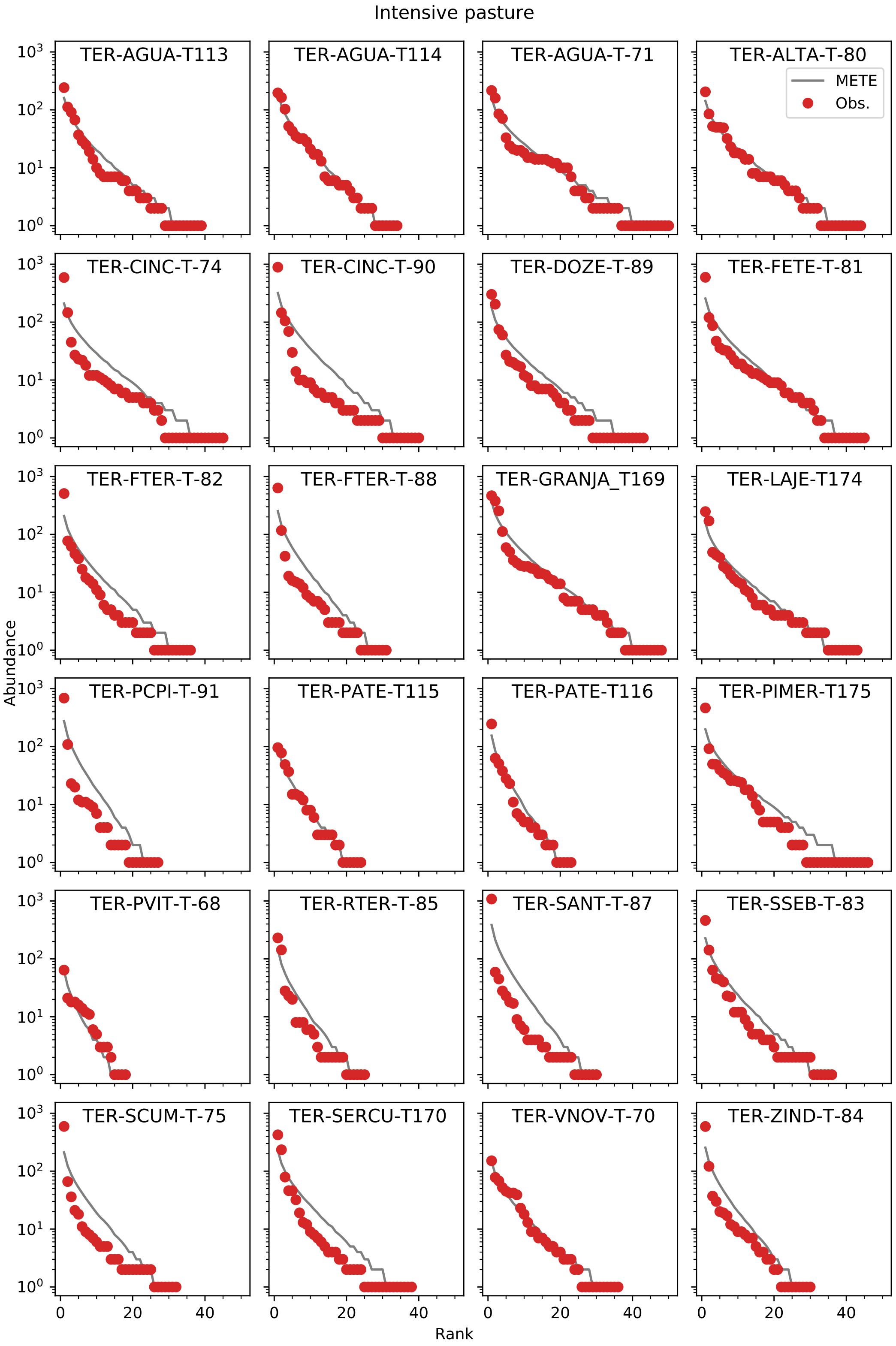


Figure S14: The rank ordered SAD at each transect in the intensive pasture.

#### Appendix S11 MRDIs at each transect

Presented in order of increasing land use intensity:

- Fig. S15 – Native forest
- Fig. S16 – Exotic forest
- Fig. S17 – Semi-natural pasture
- Fig. S18 – Intensive pasture.

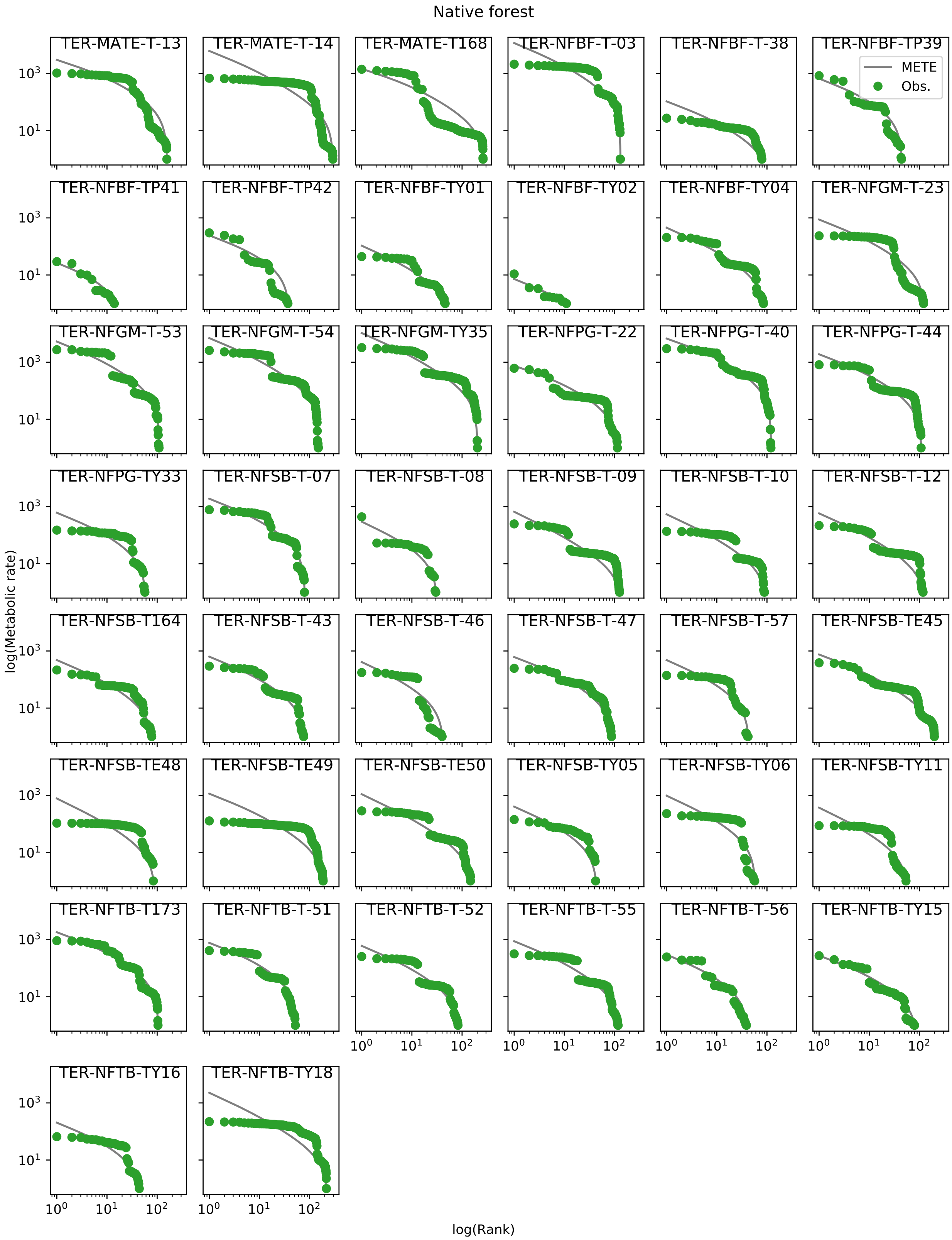


Figure S15: The rank ordered MRDI at each transect in the native forest.

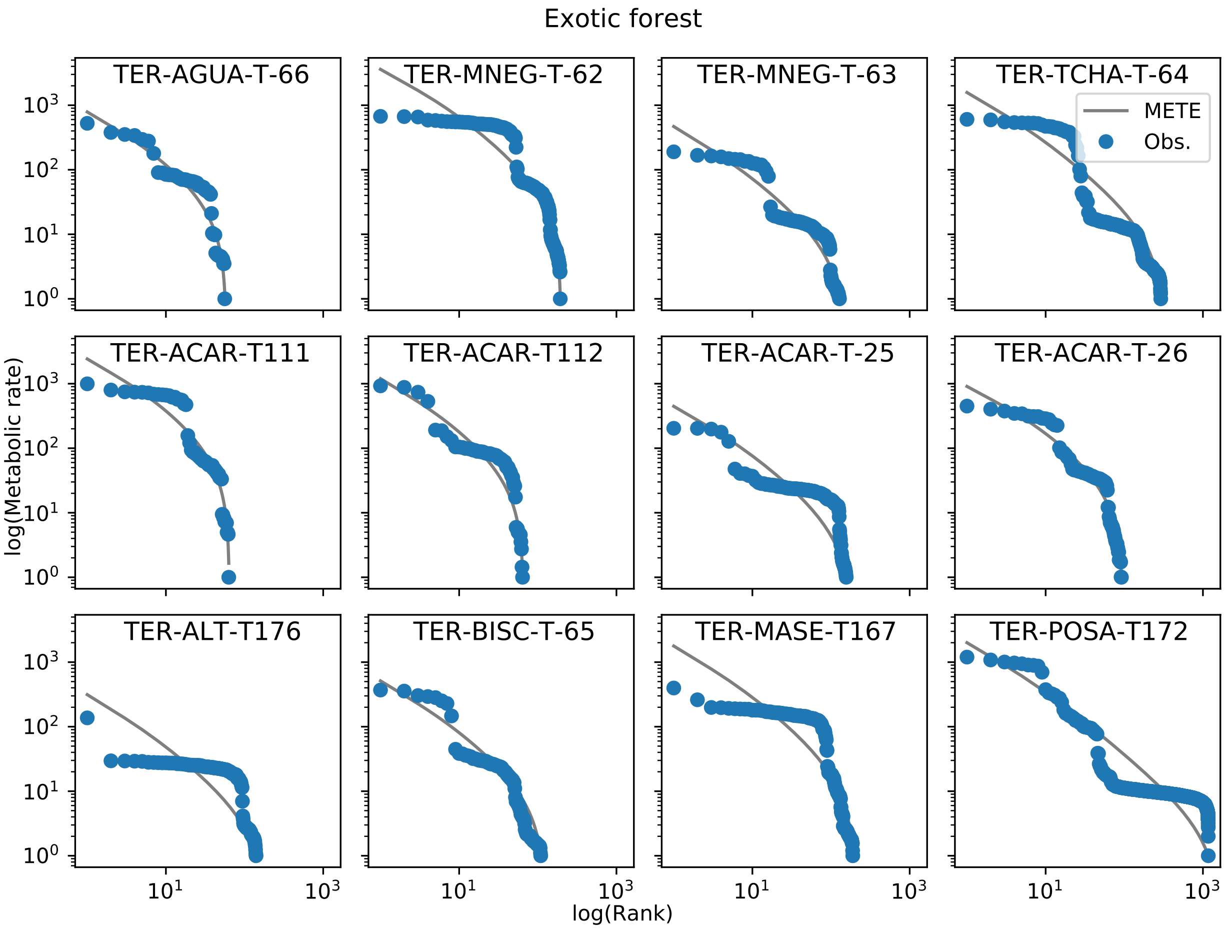


Figure S16: The rank ordered MRDI at each transect in the exotic forest.

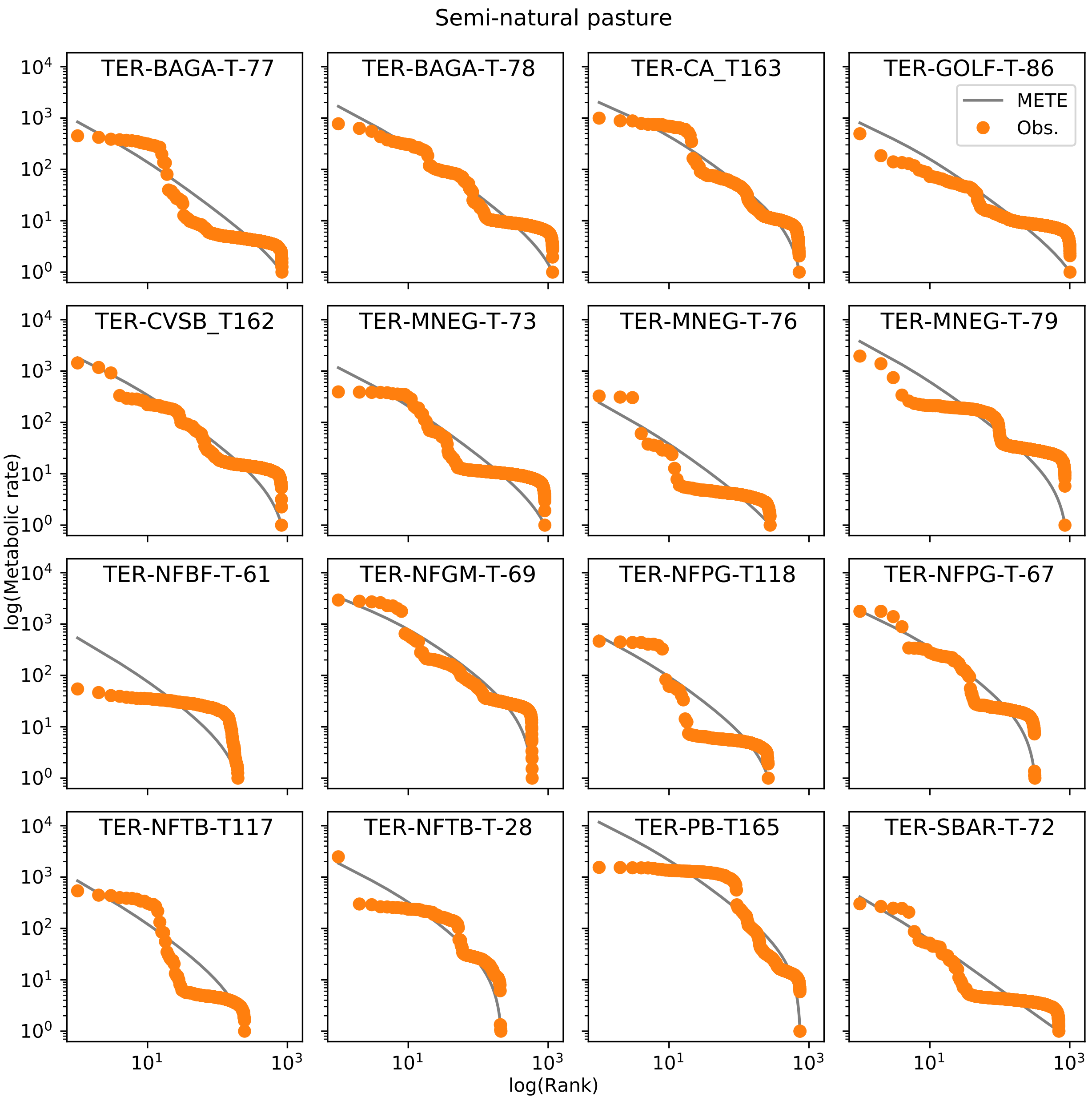


Figure S17: The rank ordered MRDI at each transect in the semi-natural pasture.

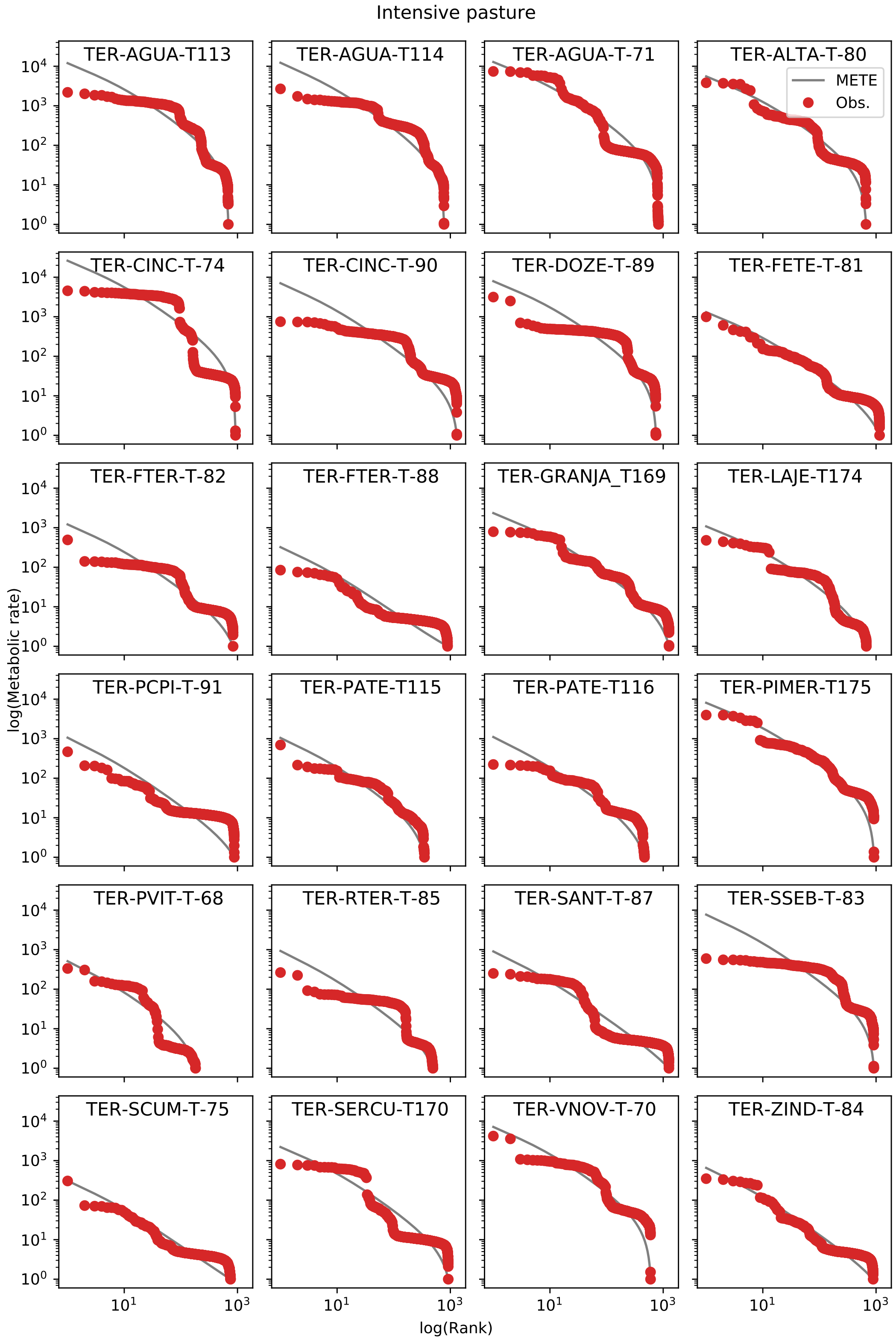


Figure S18: The rank ordered MRDI at each transect in the intensive pasture.

#### Appendix S12 SAR data on one plot

Figure S19 uses the scale-collapse of the $z - D$ relationship to display all of the data across sites together on a single plot, combining the four panels in Fig. 4.

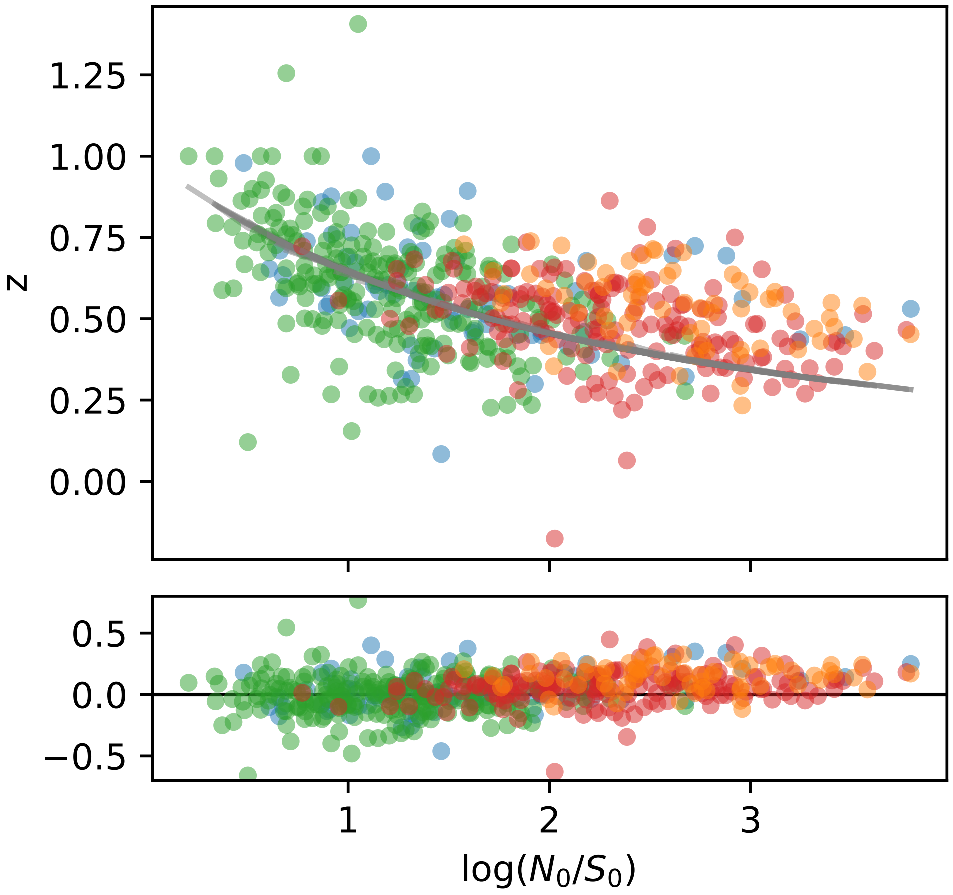


Figure S19: The species–area relationship for each transect all on a single plot, color coded by land use (colours match those in Fig. 4, with green corresponding to native forest, blue to exotic forest, yellow to semi-natural pasture, and red to intensive pasture). Each point represents a single transect at a specific scale, where the scale is determined by $\log(N_0/S_0)$. The gray lines correspond to the METE predictions. Here we have plotted the slope of the relationship on the y-axis so that all points collapse onto one universal curve. The residuals are shown below.
